# A neural entero-pancreatic pathway that regulates insulin secretion and glucose tolerance

**DOI:** 10.1101/2025.08.14.670343

**Authors:** Madina Sokolov, Alejandro Tamayo, Noah J. Levi, Randy J. Seeley, Alejandro Caicedo

**Author notes:** **Corresponding authors:** Correspondence to Madina Sokolov, Randy J. Seeley, and Alejandro Caicedo.

## Abstract

Signals from the gut enhance pancreatic secretion of insulin and thus influence glucose metabolism. This phenomenon, known as the incretin effect, is thought to be mediated by hormones secreted from enteroendocrine cells. The endocrine model, however, does not fully capture the complexity of gut-pancreas interactions. Anatomical studies identified a direct neural connection between the gut and the pancreas, known as the entero-pancreatic plexus. The role of this connection in regulating glucose metabolism remains unknown. Here we identify and functionally characterize a subpopulation of nitridergic myenteric neurons in the proximal duodenum that project directly to the pancreas. The anatomical and transcriptomic signature of these neurons places them downstream of glutamatergic and enkephalinergic enteric interneurons that indirectly respond to luminal nutrient stimuli. Their axonal projections to the pancreas contact neuro-insular ganglia and reach into the islet parenchyma. When activated chemogenetically, this circuit increases Ca^2+^ responses in pancreatic beta cells, enhances insulin secretion, and improves glucose tolerance *in vivo*. Our findings reveal a direct gut–pancreas neural pathway that complements incretin signaling in potentiating insulin secretion. This unexpectedly strong neuronal modulation of beta cell function could be harnessed to improve glycemic control in diabetes.

## Main

Blood glucose regulation depends on the capacity of pancreatic beta cells to secrete insulin in response to elevated glucose levels.^1^. In addition to this reactive response, sensory signals from the gastrointestinal tract enhance insulin secretion in anticipation of nutrient absorption.^2–5^ Such feed-forward control helps prevent glucose spikes after a meal and is often impaired in type 2 diabetes.^6,7^ Over the past decades, extensive research has focused on the roles of gut incretin hormones, vagal sensory afferents, and central neural circuits in coordinating this anticipatory glucoregulatory response.^8–16^ By contrast, the contribution of the enteric nervous system to the gut-pancreas axis remains understudied.

Enteric neurons form an extensive neural network that processes and integrates mechanical and chemical input from the gut lumen to generate output to target cells of the gut wall and viscera. Luminal cues detected by intrinsic primary afferent neurons are transmitted *via* myenteric neurons to effector cells, triggering peristaltic and secretomotor reflexes.^17–21^ Some myenteric neurons project directly to the pancreas, forming the entero-pancreatic plexus, first described by Kirchgessner and Gershon.^22–27^ A functional entero-pancreatic neural reflex has been demonstrated for the exocrine pancreas, where enteric neurons synapse onto pancreatic neurons to regulate acinar secretion in response to luminal stimuli such as glucose or mechanical stretch.^24,28–30^ This reflex engages nitric oxide and pituitary adenylate cyclase-activating polypeptide (PACAP) signaling to coordinate enzyme release with digestive demands. However, despite anatomical evidence of enteric innervation of the pancreas, a comparable reflex regulating endocrine function and hormone secretion has not been shown, and the systemic metabolic role of the entero-pancreatic plexus remains uncharacterized.

Here we investigated the direct effects of enteric neurons on insulin secretion and glucose metabolism. We combined modern tracing tools and single-nuclei transcriptomics with chemogenetic stimulation and live imaging of neuronal and beta cell function to define and characterize the phenotypes, innervation patterns, and effects of enteric neurons projecting to the pancreas. We further tested their impact on glucose metabolism in oral and intraperitoneal glucose tolerance tests. Our findings provide new insight into how enteric neurons contribute to glucose metabolism control and highlight the entero-pancreatic plexus as a novel pathway involved in regulating insulin secretion during the intestinal phase of digestion.

### Enteric neurons of the proximal GI tract project to the pancreas

Given the paucity of studies since its initial description, we first revisited the entero-pancreatic plexus using current viral approaches for anterograde and retrograde tracing. For retrograde tracing studies, we injected the recombinant adeno-associated virus (rAAV) expressing reporter mCherry under neuronal human synapsin promoter (rAAV-hSyn-mCherry, AddGene 114472) into the pancreatic duct of ChAT-GFP transgenic mice (Fig. 1a). We identified nitridergic neurons using immunohistochemistry. We inspected peripheral neuronal structures known to innervate the pancreas, namely local pancreatic ganglia, the enteric nervous system (ENS), nodose ganglia (NG) of the sensory vagus nerve, thoracic dorsal root ganglia (DRG, T9 - T11) of spinal sensory nerves, and coeliac ganglia of the sympathetic system.

**Figure 1.**
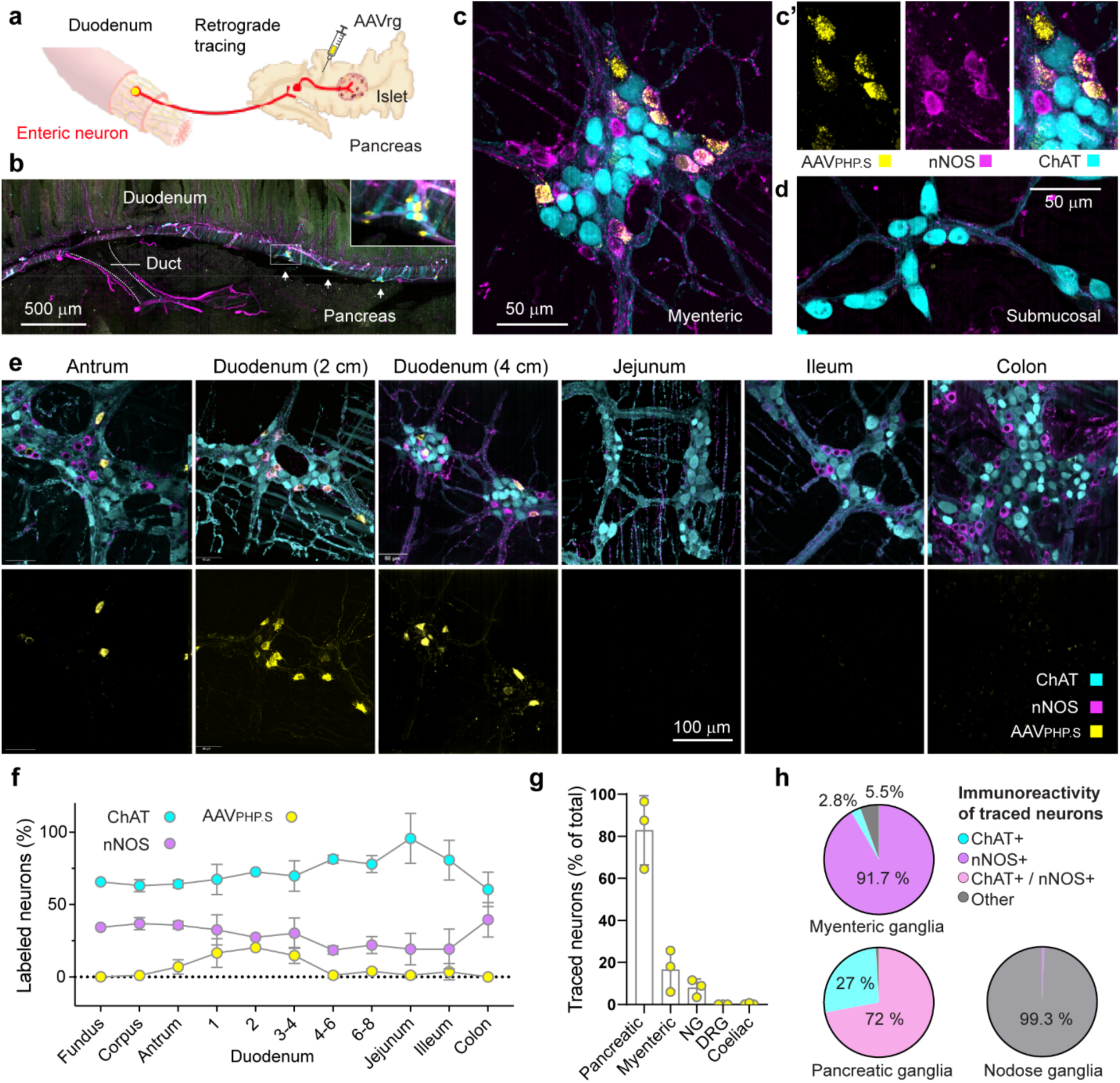
Pancreas-projecting enteric neurons are located in the myenteric plexus of the proximal gut and are predominantly nitrergic. **a**, Schematic of the entero-pancreatic neural circuit and retrograde tracing strategy. AAVrg-hSyn-mCherry was injected intraductally into the pancreas of ChAT–GFP mice. **b**, An *in toto* preparation of the duodenum and pancreas illustrating their anatomical continuity at the level of the common bile duct. Tissues were immunostained for βIII-tubulin (magenta), ChAT–GFP (cyan) and viral mCherry (yellow). **c–e**, Flat-mount preparations showing traced neurons in myenteric but not submucosal ganglia. Immunostaining for nNOS (magenta), ChAT–GFP (cyan) and mCherry (yellow). **c, c′**, Retrogradely labeled neurons in the duodenal myenteric plexus **d**, Submucosal ganglia lack labeled neurons. **e**, Distribution of traced neurons along the myenteric plexus, distances from the pylorus indicated. **f–h**, Quantification of neuronal phenotypes (3–15 randomly selected areas per region, 100–1000 neurons per area, *n* = 3 mice). **f**, Percentage of total neurons positive for ChAT, nNOS and the tracer AAV_PHP.S_. **g**, Percentages of AAV_PHP.S_+ neurons of the total neuronal population in peripheral ganglia known to innervate the pancreas: myenteric plexus (2–4 cm from pylorus), nodose ganglion (NG), thoracic dorsal root ganglia (DRG; T9–T11) and coeliac ganglia. **h**, Neurochemical identity of traced neurons in myenteric, pancreatic and nodose ganglia: ChAT+ (cyan), nNOS+ (magenta), ChAT+/nNOS+ (pink) or double-negative (grey).

Traced neurons in the duodenum were localized to the myenteric plexus, lacked Chat-GFP signal, but immunostained for neuronal nitric oxide synthase (nNOS; Fig. 1b,c). We did not find traced neurons in the submucosal plexus (Fig. 1d). In the myenteric plexus, most of the traced neurons localized to the stomach antrum and proximal duodenum within 6 cm of the pylorus (18% +/− 6), but not in more rostral or caudal regions (Figure 1c-g). The majority of traced neurons in myenteric ganglia were nitridergic (92%, Fig. 1h).

We further found traced neurons in pancreatic ganglia (81% +/− 9.5) that mainly were double-positive for nitridergic and cholinergic markers (72%; Fig. 1h). In line with previous reports^31,32^, we found traced neurons in the nodose ganglion (8% +/− 3.4), of which only 0.7% (+/− 0.5) were nitridergic (Fig. 1g,h). We could not find traced neurons in T9-T11 DRGs nor in coeliac ganglia (Fig. 1g and Extended data Fig. 2). This observation is consistent with a previous report showing that AAVrg is inefficient for tracing from the pancreas to the DRGs and coeliac ganglia.^33^ Our results show that ∼20% of myenteric neurons of the proximal duodenum project to the pancreas and indicate that these neurons are nitridergic, not cholinergic.

**Figure 2.**
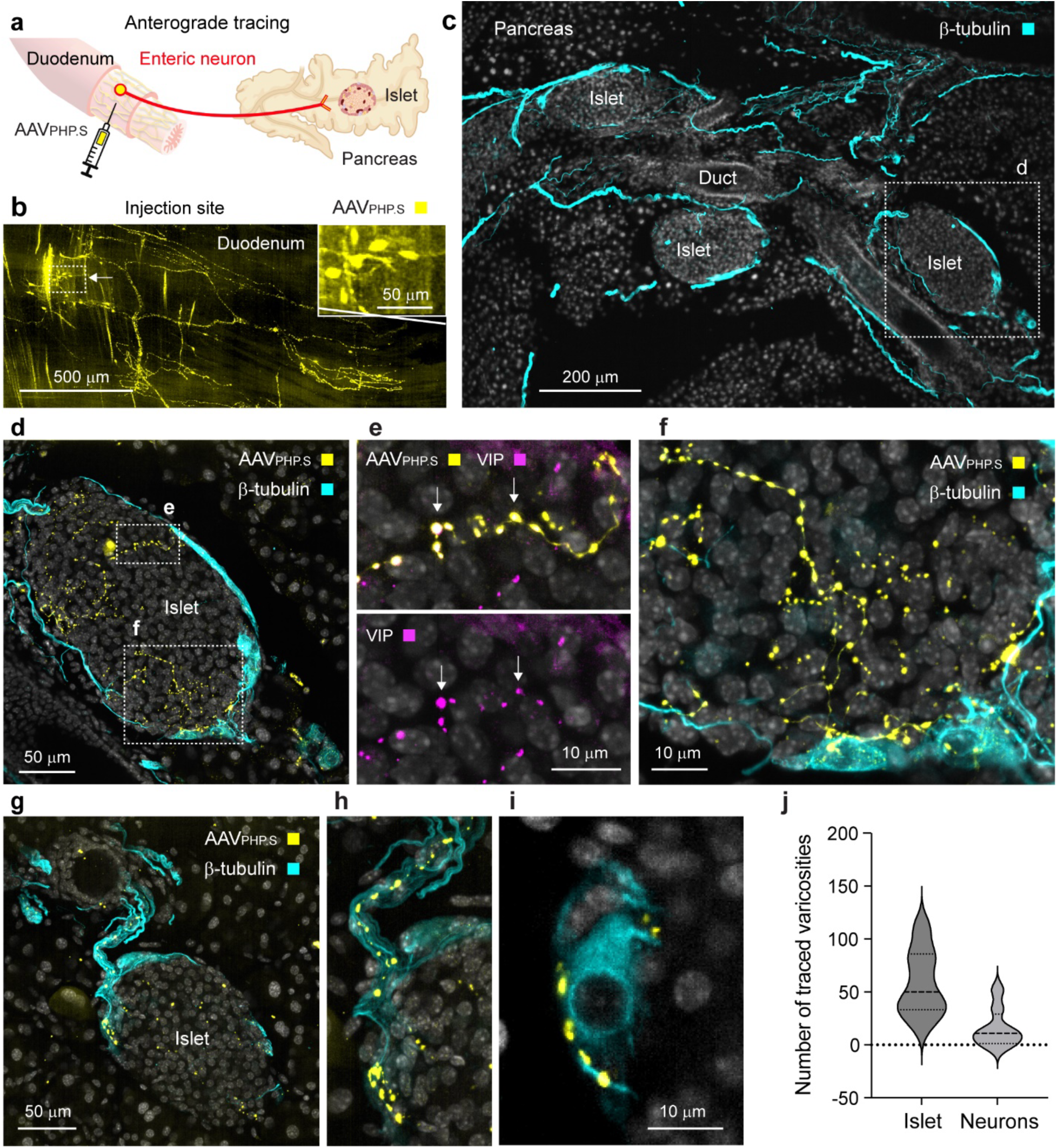
Terminals of enteric neurons in the pancreas. **a**, Schematic of the entero-pancreatic neural axis and anterograde viral tracing strategy. AAVphp.s-hSyn-mCherry was injected submuscularly into the stomach antrum and duodenum of wild-type mice. **b**, Duodenal flat mount showing a viral injection site (mCherry, yellow). **c–i**, Representative pancreatic sections immunostained for βIII-tubulin (cyan), VIP (magenta), mCherry (yellow) and DAPI (grey). **c**, Low-magnification view of a section containing three endocrine islets. **d–f**, High-magnification view of a representative islet (highlighted in c) with numerous traced varicosities in the endocrine parenchyma and fewer in a neighboring ganglion; some varicosities are VIP-positive (**e**). **g–i**, Islet with few traced varicosities in the endocrine parenchyma but more in the adjacent ganglion. **j**, Quantification of traced varicosities in endocrine islets as in **d** and **g**. Varicosities were present in 8 of 39 analyzed islets (*n* = 3 mice).

To further characterize the connection between myenteric neurons and the pancreas we performed anterograde tracing. We injected a virus of the PHP.S serotype (AAVphp.s-hSyn-mCherry) into the muscle layers of the stomach and proximal duodenum of B6 mice (Fig. 2a). Infected neurons were confined to injection sites and were not numerous (<100 neurons; Fig. 2b). We inspected pancreatic tissues for traced neuronal terminals and found terminals in 20% of pancreatic islets and islet-associated neuro-insular ganglia (Fig. 2c-I and Supplementary Video 1). The number of synaptic varicosities ranged from 20 to 111 in islets and 5 to 52 in neuro-insular ganglia (Fig. 2j). To define the phenotype of the traced fibers, we relied on immunostaining for vasoactive intestinal polypeptide (VIP) because nNOS immunostaining did not label fine neuronal terminals. We found that VIP, which is commonly expressed in nitridergic enteric neurons^34–36^, was localized in anterogradely labeled fibers (Fig. 2e). Although these qualitative results do not capture the whole extent of the entero-pancreatic innervation, they complement the retrograde tracing studies by demonstrating innervation targets within the pancreas, namely pancreatic ganglia and islets.

To control for potential viral leakage, we performed two types of sham surgeries in which the virus was applied topically to the peritoneum or into the gut lumen, simulating peritoneal or luminal leakage. We did not detect traced enteric neurons in either control group, in line with previous reports that AAV viruses require direct injection into the gut wall or retrograde transport from target tissues to infect enteric neurons^37^.

### Molecular profiling of myenteric neurons projecting to the pancreas

To investigate the molecular signatures of pancreas-projecting neurons, we acquired single-nuclei transcriptome data on enteric neurons traced from the pancreas. The dispersed nature of myenteric plexus within the gut wall makes it challenging to isolate it from the surrounding tissue. According to previous reports, enteric neurons represent around 0.004% of the total cellular makeup of the gut wall.^34^ Neurons traced from the pancreas represent under 20% of the duodenal myenteric neuronal population, which brings it to under 0.0008% of the gut wall cell population. We used INTACT (isolation of nuclei tagged in specific cell types) technique to label and enrich the subpopulation of pancreas-projecting myenteric neurons. To label nuclei, floxed Sun1-GFP mice received intrapancreatic injection of AAVrg driving the expression of Cre-recombinase under the synapsin promoter (AAVrg-hSyn-Cre), analogous to the retrograde tracing technique described earlier. Nuclear GFP labeling matched the cytoplasmic retrograde tracing described earlier in that it was confined to nitridergic neurons of the myenteric plexus (Figs. 1 and 3). Nuclei were isolated from snap-frozen duodenal samples, and GFP+ nuclei were enriched by FACS sorting and sequenced on a 10x platform. Our approach yielded high-quality transcriptomic data from 111 GFP+ enteric neurons.

**Figure 3.**
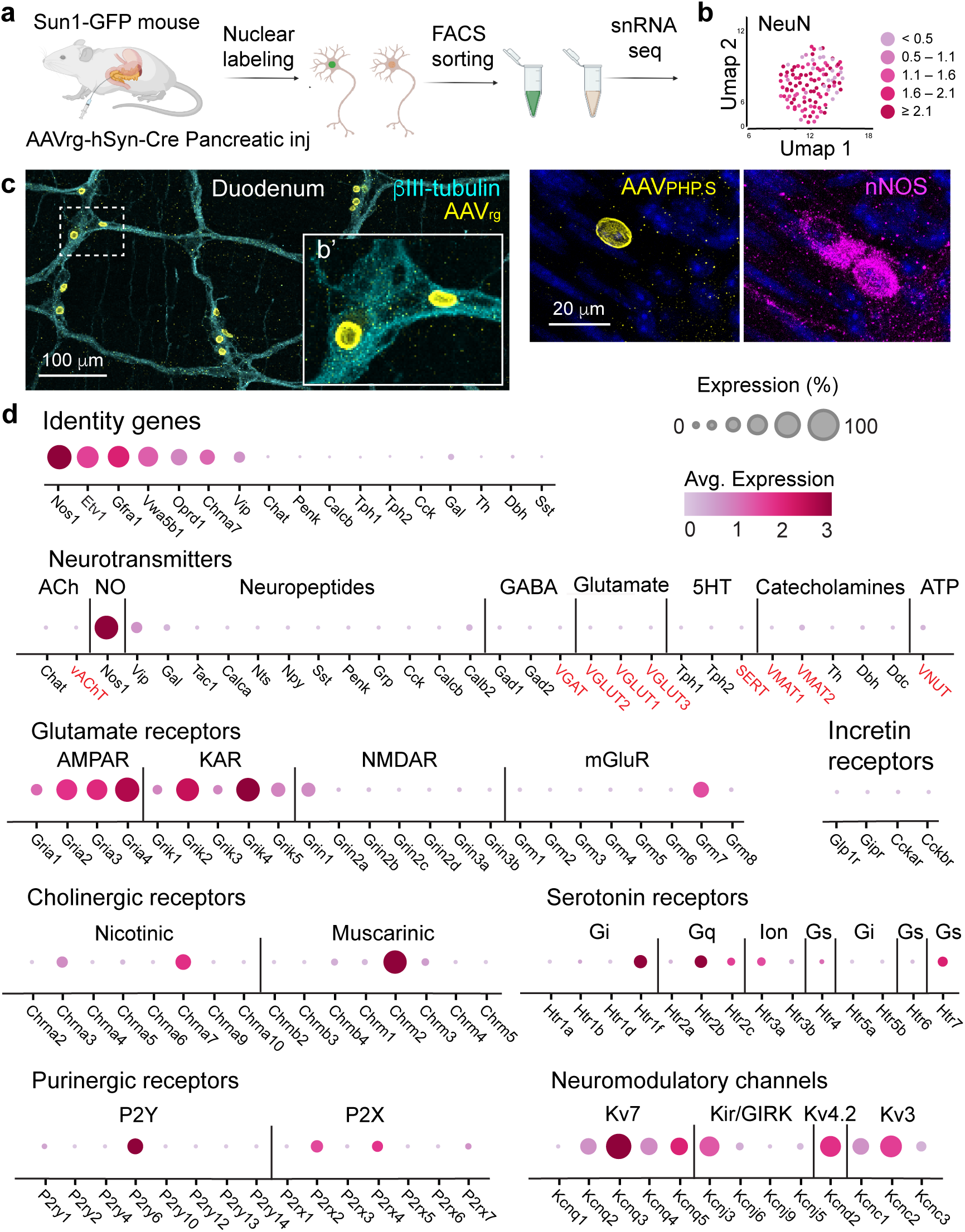
Molecular profiling of entero-pancreatic neurons reveals nitridergic secretory and glutamate receptor signatures. **a,** Schematic of the experimental design where entero-pancreatic neurons were labeled with GFP using the INTACT technique followed by snRNA-seq. To achieve population-specific nuclear labeling, floxed Sun1-GFP transgenic mice received intraductal pancreatic injection of AAVrg-hSyn-Cre. **b,** Reference UMAP of 111 GFP+ enteric nuclei from mouse duodenum. **c,** Representative images of the duodenal flat mounts depicting Sun-1-GFP+ nuclei in nNos+ myenteric neurons. Immunostaining for bIII-Tubulin (cyan), nNos (magenta), vAChT (white), DAPI (blue), and endogenous Sun1-GFP (yellow). **d,** Transcriptomic signature of pancreas-projecting enteric neurons reported in a series of dot plots for genes associated with major groups of neurotransmitters and neuronal receptors. Dot size reflects the fraction of nuclei expressing the gene, dot color indicates the mean expression level in expressing nuclei. Some of the gene names were swapped with the names of the protein they encode, for ease of identification (red). For a comparison of the transcriptional profiles of entero-pancreatic neurons with all enteric neurons from Drokholyanski *et al.*, see Extended Data Fig.3.

Our analyses revealed a homogeneous nitridergic neuronal population with Nos1 being the most highly expressed gene (Fig. 3). A proportion of these neurons also expressed VIP, but none expressed other signaling neuropeptides (Fig. 3d). They did not express genes required for the molecular machinery that define cholinergic, GABAergic, serotonergic, catecholaminergic, and glutamatergic neurons (Fig. 3d). We examined receptor expression and identified high expression levels for all four AMPA receptor subunits and the kainate receptor subunits Grik2 and Grik4, indicating that these neurons are fully equipped to respond to excitatory glutamatergic input. High levels of delta opioid receptor Oprd1 indicates inhibitory modulation by endogenous enkephalins. High expression levels for the muscarinic cholinergic receptor Chrm2, known to couple to the G protein Gαi, suggests that cholinergic input to these neurons is inhibitory. Moderate expression of serotonergic and purinergic receptors indicates that serotonin and ATP may modulate these neurons. Of the ion channels known to control neuronal excitability, voltage-gated potassium channels Kv7, GiRK, Kv4.2, and Kv3 were abundantly expressed in most neurons.

We compared the transcriptomic profile of entero-pancreatic neurons to the mouse small intestine dataset from Drokhlyansky et al.^34^ and found a high degree of overlap with the PIMN1 cluster, corresponding to Primary Intrinsic Motor Neurons 1 (Extended Data Fig. 3). This cluster is defined by a unique gene signature that includes *Nos1*, *Etv1*, *Gfra1*, *Vwa5b1*, *Oprd1*, *Chrna7*, and *Vip*, and notably lacks expression of neuropeptides and neurotransmitter enzymes common to other ENS neuronal subtypes, such as *Chat, Penk, Calcb, Tph1, Tph2, Cck, Gal, Th, Dbh*, and *Sst* (Extended Data Fig. 3).

Our findings that myenteric neurons projecting to the pancreas are uniformly nitridergic, but not cholinergic or serotonergic, contrast with early anatomical reports from Kirchgessner and Gershon.^22,24,25,27^ However, our results are consistent with their work on entero-pancreatic reflex controlling acinar tissue^29,30^ and align with our own tracing studies.

### Entero-pancreatic neurons influence pancreatic beta cell function

To investigate the effects of the entero-pancreatic neural plexus on islet physiology and glucose metabolism, we used a chemogenetic approach to modulate activity of these neurons. We injected AAV expressing Cre-dependent designer receptor activated by designer drug (DREADD) - hM3D(Gq) (AAVrg-hSyn-DIO-hM3D(Gq)-mCherry) or control AAV (AAVrg-hSyn-DIO-mCherry) into the pancreas of nNos-Cre mice. This approach enabled us to selectively target populations of nitridergic neurons projecting to the pancreas (Fig. 4). Out of the peripheral neural ganglia projecting to the pancreas (ENS, NG, DRG, coeliac) only entero-pancreatic neurons showed DREADD expression, in line with our data showing only myenteric ganglia contained pancreas-projecting neurons that were nitridergic (Figs. 1g-h and 4b). To verify that intraperitoneal administration of DREADD agonist clozapine nitric oxide (CNO) activates entero-pancreatic neurons, we adapted an intravital imaging approach^38^ to record Ca²⁺ responses in DREADD-expressing myenteric neurons *in vivo* (Fig. 4f). In contrast, experiments on living pancreatic slices showed that bath-applied CNO failed to elicit Ca²⁺ responses in local pancreatic neurons (Fig. 4g).

**Figure 4.**
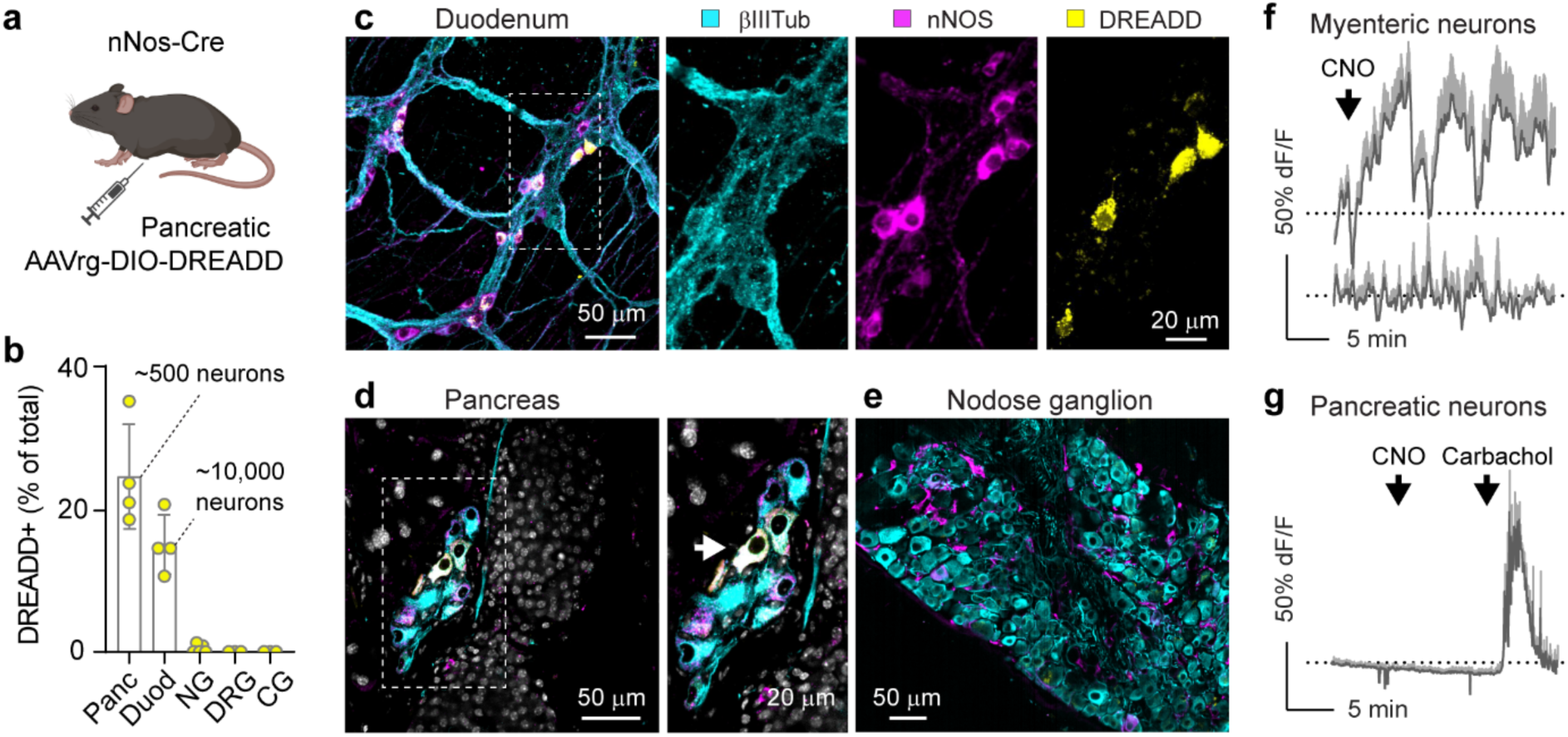
Chemogenetic manipulation of entero-pancreatic nitrergic neurons. **a,** Schematic of the experimental design. *nNos*-Cre mice received intraductal pancreatic injection of AAVrg-DIO-hM3D(Gq)-mCherry to express the excitatory DREADD receptor in entero-pancreatic neurons. **b,** Quantification of DREADD+ neurons in peripheral neural ganglia. Pancreatic ganglia (Panc), myenteric plexus (Duod, 2–4 cm from pylorus), nodose ganglion (NG), thoracic dorsal root ganglia (DRG; T9–T11), and coeliac ganglia (CG). Counts were based on 3–15 randomly selected areas per region (Pancreas: 408 neurons from 34 ganglia; Duodenum: 737 neurons from 34 ganglia; *n* = 4 mice). No mCherry signal was detected in NG, DRG, or CG. Neuron counts were normalized to total tubulin-positive neurons. Numbers above bars indicate estimated total neuron numbers targeted by this approach (∼500 pancreatic and ∼10,000 myenteric neurons; see Methods for details). **c–e,** Representative images of a duodenal flat mount (c) and sections of pancreatic and nodose ganglia (d, e) showing co-localization of DREADD-mCherry⁺ and nNos⁺ neurons. Immunostaining: βIII-tubulin (cyan), nNos (magenta), vAChT (white), DAPI (blue), endogenous Sun1-GFP (yellow). **f,** *In vivo* Ca^2+^ imaging traces from nitrergic myenteric neurons showing responses (changes in mean fluorescence intensity, ΔF/F₀) recorded using intravital imaging. CNO was administered intraperitoneally (5 mg/kg). Top: responding neurons; bottom: non-responding neurons. **g,** *In vitro* Ca^2+^ imaging traces from pancreatic neurons showing responses (ΔF/F₀) recorded *ex vivo* in living pancreatic slices. CNO (20 μM, bath-applied) did not elicit Ca^2+^ responses but the control stimulus carbachol (30 μM) did.

To study the effect of chemogenetic activation of entero-pancreatic neurons on pancreatic beta cells, we performed intravital imaging of the pancreas to record Ca^2+^ responses in beta cells *in vivo*. Mice that expressed DREADD in pancreas-projecting enteric neurons received intraperitoneal injections of an AAV8-Ins-GCaMP6 virus driving expression of the Ca^2+^ indicator GCaMP6 under the insulin promoter (Fig. 5a). In these mice, we exposed the head of the pancreas, stabilized it in a vacuum imaging chamber, and imaged it under a confocal microscope (Fig. 5b and Supplementary Video 2). Isoflurane anesthesia induced hyperglycemia in a range of 250 – 400 mg/dl. This, however, was not due to decreased beta cell activity and insulin secretion because beta cells showed strong, persistent Ca^2+^ oscillations with periods ranging from 2-5 min in pancreatic islets. Thus, under this anesthesia, islets were working at high capacity. Upon CNO injection (5 mg/kg), the width of the Ca^2+^ peaks increased, with maximum peaks appearing 7 - 10 min post CNO injection (Fig. 5c, d; Extended data Fig. 4-7; Supplementary Video 2).

**Figure 5.**
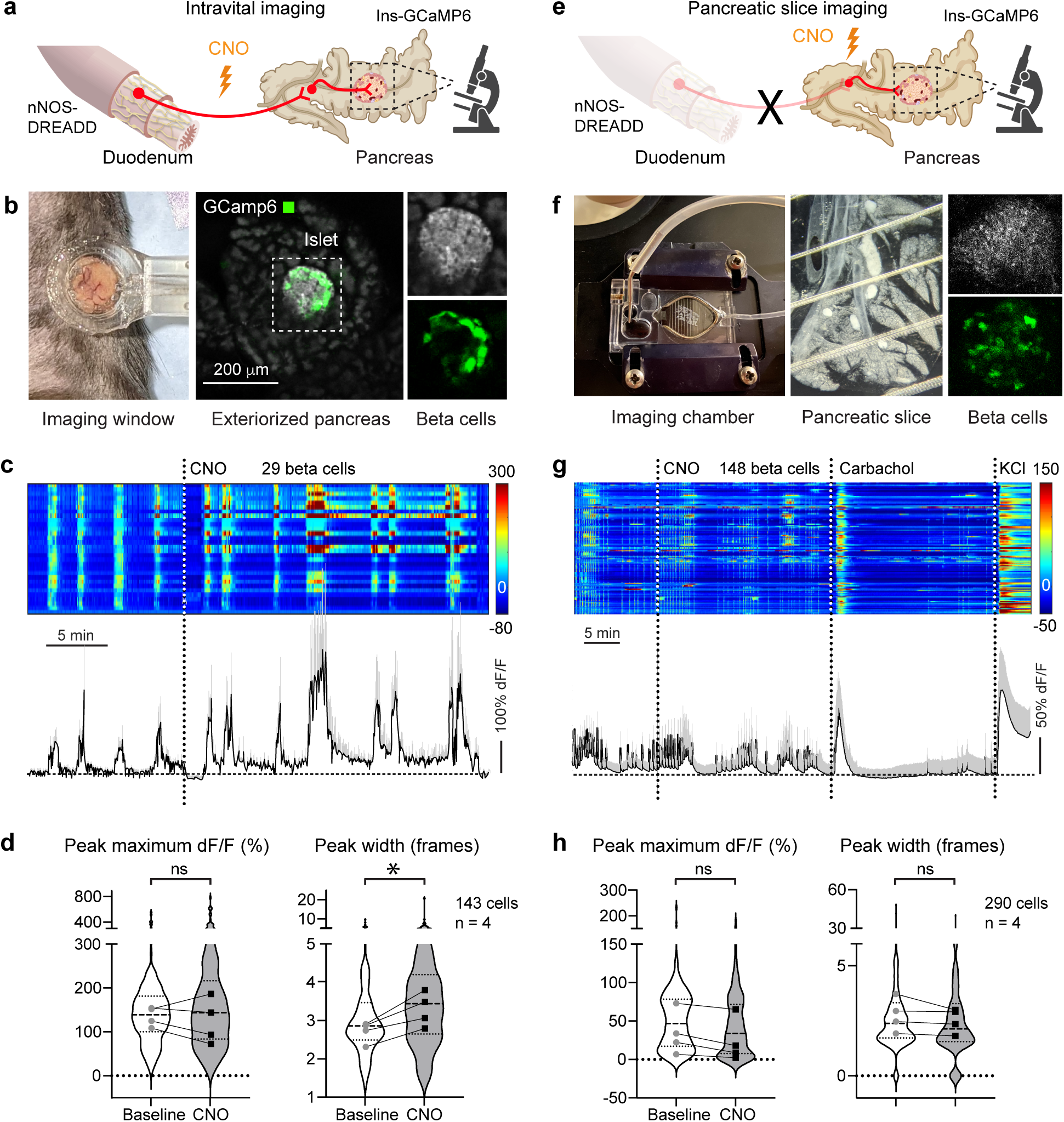
Chemogenetic stimulation of entero-pancreatic neurons enhances pulsatile Ca^2+^ responses in pancreatic beta cells *in vivo* with an intact gut-pancreas connection, but not in living pancreatic slices *ex vivo*. *nNos*-Cre mice received intraductal pancreatic injections of AAVrg-DIO-hM3D(Gq)-mCherry to express the excitatory DREADD receptor in entero-pancreatic neurons and IP injections of AAV8-Ins-GCaMP6 to drive expression of Ca^2+^ indicator in pancreatic beta cells. **a–b,** Illustration of intravital imaging of pancreatic beta cells with a preserved gut–pancreas anatomical connection. The pancreas was imaged *in vivo* through an intravital imaging window. Islets were identified by insulin backscatter (gray) and GCaMP6-positive beta cells. **c-d,** Representative heat maps and an average trace showing *in vivo* Ca^2+^ responses in beta cells. Horizontal rows in the heatmap represent individual beta cells, color intensity corresponds to the change in the mean fluorescence intensity (ΔF/F₀). *In vivo* imaging was performed under isoflurane anesthesia. Anesthesia increased glycemia to 250-400 mg/dl, which elicited pulsatile, synchronized Ca^2+^ responses in beta cells. Injecting CNO (5 mg/kg, ip) amplified these responses (quantifications are shown in **d**; 143 beta cells from n = 4 mice). **e–f,** Illustration of imaging of beta cell function in living pancreatic slices. Islets were identified by insulin backscatter and GCaMP6-positive beta cells. Imaging was performed in slices continuously exposed to a glucose concentration (7mM) that stimulates mouse beta cells at a low level, as seen by the Ca^2+^ oscillations. **g-h,** Representative heat maps and an average trace showing *ex vivo* Ca^2+^ responses in beta cells in slices. Horizontal rows in the heatmap represent individual beta cells, color intensity corresponds to the change in the mean fluorescence intensity (ΔF/F₀). CNO (20 mM), carbachol (30 mM), and KCl (25mM) were applied via a perfusion system for 2 min each. Carbachol and KCl depolarizations elicited strong Ca^2+^ responses in beta cells, but CNO did not. Note that pulsatile activity was not affected by CNO (quantifications are shown in **h**; 424 beta cells from n = 4 mice) For both *in vivo* and *ex vivo* imaging we calculated Ca^2+^ peak maxima and peak width of the pulses after CNO application **(d** and **h**). Data are mean ± s.e.m. *P* < 0.05; paired *t*-test. The image used in **f** to illustrate the pancreatic slice was adapted from Qureshi et al., 2023.^61^

To validate that these effects depended on an intact gut-pancreas axis and were not contributed by nitridergic neurons in pancreatic ganglia (Fig. 4b,d), we prepared living pancreatic slices from the same animals after completion of the intravital imaging for *in situ* imaging (Fig. 5e,f). Beta cells responded to high glucose concentration (7 mM) with Ca^2+^ oscillations, but these responses were not affected by exposure to CNO (Fig. 5 g,h and Supplementary Video 3). That is, although intraductal injection of AAVrg-hSyn-DIO-hM3D(Gq) induced DREADD expression in some nitridergic pancreatic neurons (Fig.4b,d), stimulation of the pancreas with CNO did not elicit a response in local neurons and did not affect beta cell activity. This contrasts with our results showing that chemogenetic stimulation of cholinergic intrapancreatic neurons readily potentiated beta cell responses to high glucose in living pancreatic slices.^39^ These control experiments let us conclude that acute *in vivo* chemogenetic activation of nitridergic entero-pancreatic neurons enhanced pancreatic beta cell activity under stimulatory glycemic conditions. Our results thus establish that the connection between myenteric neurons and their pancreatic targets is not only anatomical but also functional.

### Entero-pancreatic neurons modulate glucose metabolism

We investigated the effects of chemogenetic activation of nitridergic entero-pancreatic neurons on glucose metabolism. These experiments were performed on the same animals prior to intravital Ca^2+^ imaging (Fig 6a,b). Our strategy consisted of comparing the same cohort of mice before and after intrapancreatic injection of either DREADD (AAVrg-hSyn-DIO-hM3D(Gq)-mCherry) or control (AAVrg-hSyn-DIO-mCherry) viruses. After 4 weeks of post-surgical recovery, the same mice underwent intraperitoneal and oral glucose tolerance tests (IPGTTs and OGTTs) after injection of CNO (5 mg/kg) or saline. We thus compared intraperitoneal and oral glucose tolerance in the same animals before and after DREADD infection in the presence and absence of CNO. Of note, the OGTTs include an intestinal contribution to insulin secretion, while in the IPGGTs beta cells are only stimulated by circulating glucose levels.

**Figure 6.**
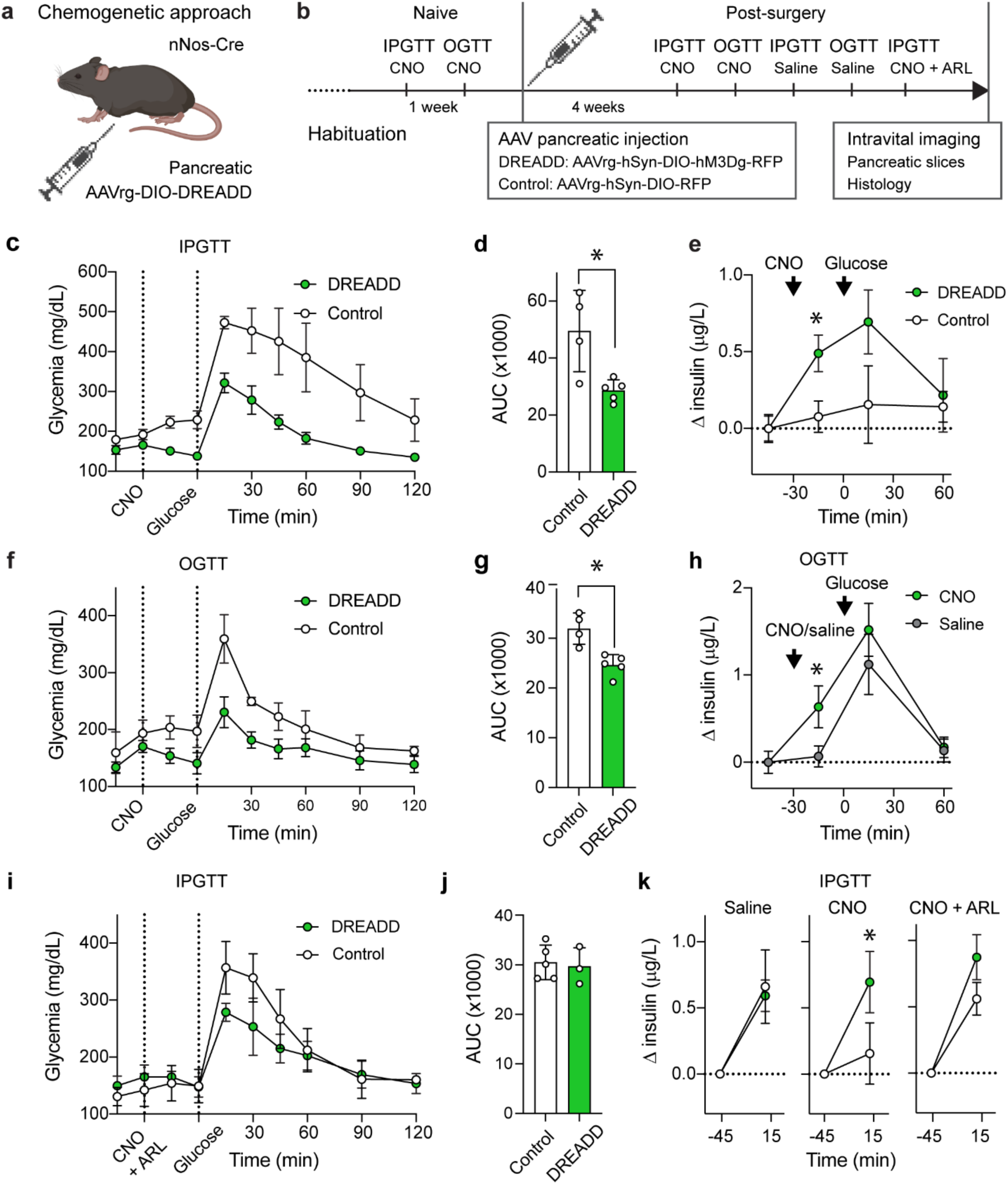
Chemogenetic activation of entero-pancreatic neurons improves glucose tolerance and enhances insulin secretion. **a, b,** Schematic of the experimental design: *nNos*-Cre mice received intraductal pancreatic injection of AAVrg-DIO-hM3D(Gq)-mCherry or control virus. The timeline indicates IPGTTs and OGTTs performed before and after viral injection. **c–e,** IPGTT following CNO injection (5 mg/kg, ip, 30 min prior to glucose). Glucose (c) and insulin levels (e) in DREADD-expressing (n = 5) and control mice (n = 4). (d) Area under the curve (AUC) for glucose in (c). **f, g,** OGTT following CNO injection. Glucose levels (f) and AUC (g) in DREADD and control mice. **h,** Insulin levels during OGTT in DREADD-expressing mice following CNO or saline injection. **i–k,** IPGTT with co-injection of CNO and nNOS inhibitor ARL17477 (10 mg/kg). Glucose (i), insulin (k), and AUC (j) in DREADD and control mice. Data are mean ± s.e.m. *P* < 0.05; unpaired two-tailed *t*-test for bar graphs; two-way ANOVA with multiple comparisons for insulin time course.

CNO injection led to immediate drops in glycemia and increases in insulin secretion in DREADD-expressing animals before they were challenged with glucose in the IPGTTs and OGTTs (Fig. 6c,e,f,h and Extended Data Figs. 8, 9). In these mice, CNO injection improved glucose tolerance in IPGTTs and OGTTs (Fig. 6c,d,f,g and Extended Data Figs. 8, 9). The glucose excursion of the IPGTTs in mice injected with CNO was comparable to the glucose excursion of the OGTT in control animals. Insulin secretion at 15 min post glucose administration was significantly higher in IPGTT but not OGTT. CNO administration did not affect glucose tolerance in the control group that did not express DREADD and naive animals prior to the virus injection (Extended Data Fig. 8). Likewise, saline did not affect glucose tolerance and insulin secretion in DREADD-expressing animals (Fig. 6k and Extended Data Figs. 8, 9). These data indicate that acute activation of entero-pancreatic neurons improves glucose tolerance and enhances insulin secretion.

To determine the role of nitridergic signaling in entero-pancreatic neurotransmission, we conducted an IPGTT in the presence of the nNOS inhibitor ARL17477 (10 mg/kg; Fig. 6i,j). This treatment abolished the difference in glucose excursion and insulin secretion seen after CNO injection in mice expressing DREADD in entero-pancreatic neurons (compare Fig. 6d-e with Fig. 6i-k).

## Discussion

Despite significant advances in understanding neuronal control of metabolism^15,40,41^, the role of the enteric nervous system in metabolic processes remains largely unexplored. Earlier studies by Kirchgessner and Gershon revealed a functional neuronal connection between the ENS and the pancreas.^22–25,27,30^ Although highlighted in reviews^20,40,42^, these findings have not been independently validated. Our study confirms the direct entero-pancreatic neural connection. Additionally, we find that pancreas projecting neurons (a) are nitridergic myenteric neurons, (b) innervate pancreatic islets and neuro-insular ganglia, (c) modulate beta cell activity, and (d) impact glucose metabolism.

The transcriptomic profile of entero-pancreatic neurons reported here offers a glimpse of their identity within the ENS circuitry. Entero-pancreatic neurons express genes of inhibitory motor neurons such as *Nos1*, *Gfra1*, *Etv1*, *Oprd1*, *Chrna7*, *Grm7*, and *Dgkb* (classified as the PIMN1 subset of putative inhibitory motor neurons in Drokholyanski et al. (Extended Data Fig. 3).^34^ Our data further show that entero-pancreatic neurons express receptors for glutamate (*Gria2-4*, *Grik2*, *Grik4*, *Grm7*), acetylcholine (*Chrna7*, *Chrm2*), and enkephalin (*Oprd1*), suggesting that they are potentially activated by glutamatergic interneurons that can co-release acetylcholine with glutamate.^21^ These glutamatergic interneurons, most likely of the putative interneuron 1 (PIN1) subset (Extended Data Figure 3)^34^, were recently shown to contact inhibitory nitridergic motor neurons.^21^ These PIN1 interneurons, in their turn, likely receive input through nicotinic transmission from myenteric sensory neurons of the PSN1 subset (Extended Data Fig. 3).^18,34^ A recent study demonstrated that these PSN1 neurons are key sites of initial luminal glucose signal processing and integration within the ENS.^18^ Thus, we propose a model where luminal glucose is first detected at the mucosal surface by enteroendocrine cells that release serotonin to activate 5-HT_3_R-expressing sensory nerve endings of myenteric PSN1 neurons^18^, which then release acetylcholine to activate glutamatergic interneurons innervating entero-pancreatic neurons (Fig. 7 and Extended Data Fig. 3).

**Figure 7.**
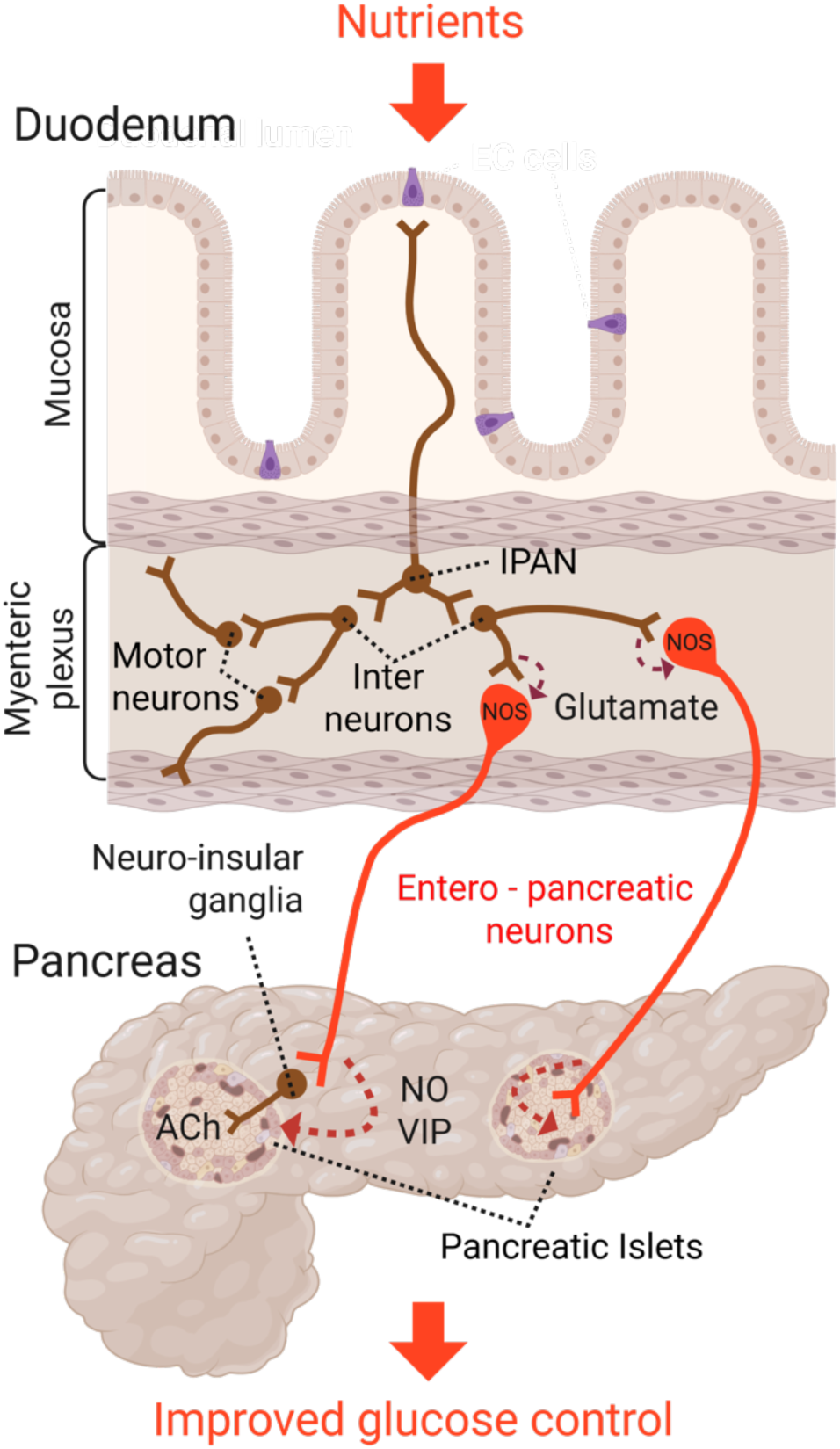
Simplified schematic of proposed model for the entero-pancreatic circuit regulating insulin secretion and glucose metabolism. Luminal glucose is first detected at the mucosal surface by enteroendocrine cells (purple) that release serotonin to activate 5-HT_3_R-expressing sensory nerve endings of myenteric IPAN sensory neurons, which then release acetylcholine to activate glutamatergic interneurons innervating entero-pancreatic neurons. These neurons project to the pancreas, where they innervate local neuronal ganglia and pancreatic islets. When activated, entero-pancreatic neurons stimulate beta cells, thus potentiating insulin secretion and improving glycemic control.

Our molecular profiling further shows *Nos1* as the most prominently expressed gene in entero-pancreatic neurons, with lower levels of expression of VIP in a subset of cells. These neurons displayed minimal levels of cholinergic, GABAergic, and glutamatergic markers or other signaling neuropeptides. Nitridergic motor neurons in the enteric nervous system produce and release NO to facilitate peristaltic muscle relaxation. The role of NO in pancreatic islet physiology, however, remains speculative. While excessive NO production in pathological conditions can cause beta cell dysfunction and suppress insulin secretion, low concentrations of NO have been found to enhance insulin secretion in response to glucose by promoting Ca^2+^ release from mitochondria.^43–49^ NO is known to produce major vascular effects.^50^ NO released from nerve endings could influence blood flow in the pancreatic islet, thus affecting insulin secretion and glucose homeostasis.^51^ How entero-pancreatic neurons release NO and VIP to modulate insulin secretion from beta cells remains to be determined.

Our findings demonstrate that *in vivo* chemogenetic activation of nitridergic entero-pancreatic neurons significantly increases beta cell activity under hyperglycemic conditions. This effect was absent in pancreatic slices lacking the gut-pancreas connection. Activating this neural pathway in awake mice rapidly reduced blood glucose levels and increased insulin secretion. This effect was also evident during both intraperitoneal and oral glucose tolerance tests, where glucose tolerance improved due to notable, significant increases in insulin secretion. Although we stimulated this neural pathway artificially with chemogenetic tools, luminal nutrient signals likely activate this pathway, too. A recent study provided strong evidence that enteric neurons respond to different luminal chemicals, including glucose.^18^ These sensory neurons are connected to downstream interneurons and motor neurons that include the entero-pancreatic population, as described above. We therefore conclude that sensory cues from the intestinal epithelium not only elicit secretion of incretin hormones but also trigger a faster neuronal response that is relayed to pancreatic beta cells during the intestinal phase of digestion. By directly targeting pancreatic islets, the entero-pancreatic neural axis adds speed and precision to the anticipatory regulation nutrients in the gut lumen exert to contribute to overall glucose homeostasis.

Patients with type 2 diabetes (T2D) frequently experience peripheral neuropathies.^52^ Neuropathy of nitrergic neurons is recognized as a contributor to gastroparesis in the later stages of the disease.^53^ Although the status of nitrergic enteric innervation in the early stages of T2D remains unclear, animal studies have shown that NOS levels are significantly reduced even in the initial phases of diabetes progression.^54,55^ Given our findings on the role of nitridergic enteric neurons in glucose metabolism, it is likely that early changes in NO production may play a role in the pathogenesis of diabetes. This underscores the importance of further research to understand better how alterations in nitrergic signaling could contribute to both gastrointestinal and metabolic dysfunctions associated with T2D.

## Methods

### Mouse Models

Experiments were conducted according to protocols and guidelines approved by the Institutional Animal Care and Use Committees of the University of Miami and the University of Michigan. All experiments used adult males aged 8–16 weeks. Mice were maintained on a 12-h light/dark cycle, and provided standard chow and water ad libitum.

For retrograde tracing studies we used Chat-eGFP mice (Jax 007902). For anterograde tracing we used C57/B6 mice (Jax 000664). For sequencing experiment, we used CAG-LSL-Sun1-sfGFP mice (Jax 021039). For Chemogenetic experiments we used nNos-Cre mice (Jax 017526).

### Surgical procedures

All surgical procedures were performed under isoflurane anesthesia (1,5 – 2 %). Animals received a single injection of Buprenorphine extended release to alleviate post-surgical pain and discomfort.

### Pancreatic injections

Retrograde AAV viruses (AAVrg, titer ∼ 10^11^ vg/ml) were delivered to the pancreas via intraductal infusion as previously described.^31,56^ Briefly, AAVrg was infused into the pancreas through the common bile duct at a rate of 7 μl min⁻¹ until a total volume of 7 - 10 μl per gram of body weight was administered. For retrograde tracing, three ChAT-eGFP mice received injection of the AAVrg-hSyn-mCherry (AddGene 114472), two control mice received either IP or intraluminal injection of the same virus. For sequencing experiment, six CAG-LSL-Sun1-sfGFP mice received injection of AAVrg-hSyn-Cre-P2A-dTomato (AddGene 107738), tissue from five mice were used for histology, tissue from one mouse was used for sequencing. For chemogenetic experiments, five nNos-Cre mice received injection of AAVrg-hSyn-DIO-hM3D(Gq)-mCherry (AddGene 44361), and five mice received injection of the control virus AAVrd-hSyn-DIO-mCherry. We allowed 4 weeks for the proper retrograde transport of the virus.

### Gut injections

Peripheral nervous system - specific AAV virus (AAV-PHP.S-CAG-tdTomato, Addgene 59462) was injected submuscularly into the stomach antrum and proximal duodenum of six mice as previously described.^57^ Briefly, Viral constructs (CAV2-Cre-GFP or AAVrg-pmSyn1-EBFP-Cre) were loaded into a Nanofil™ 10 μl syringe with a 36G needle and Silflex tubing (World Precision Instruments), mounted on a pedal-operated syringe pump. Multiple 0.05 ml injections were delivered into submucosal punctures across the stomach antrum and proximal duodenum (Extended Data Fig. 1). Four control mice underwent controlled surgery where the same virus was applied topically to the intestines. We allowed 4 weeks for the proper anterograde transport of the virus.

### Single-nucleus RNA-seq

Nuclear labeling of enteric neurons projecting to the pancreas was achieved by injecting AAVrg-hSyn-Cre-P2A-dTomato virus (AddGene 107738) into the pancreas of CAG-LSL-Sun1-sfGFP mice (Jax 021039). 4 weeks post-surgery, duodenum (first 5 cm from pylorus) was dissected out, flashed with PBS to remove luminal contents, snap frozen, and stored at −80C. Tissue was homogenized with dounce homogenization as previously described (PMID: 32888429). Nuclei were isolated using the Nuclei Pure Prep Kit (Sigma-Aldrich, NUC-101). Lysis buffer (LB) was supplemented with 80 U/ml RNase Inhibitor (Sigma-Aldrich, 3335399001). All steps were performed on ice.

Frozen samples were homogenized in LB using Dounce homogenizer until they were fully dissociated. Homogenate was filtered through 30 μm MAC strainer and centrifuged at 500 rcf for 5 min at 4C. Filtration and centrifugation step was performed twice. Resultant nuclear pellet was resuspended in wash buffer. Nuclei were labeled with propidium iodide (PI, 10 ug/ml) and FACS sorted for GFP⁺/PI⁺ population using a MoFlo Astrios Cell Sorter. Sorting gates were set based on unstained controls, single-color controls, and forward/side scatter to exclude debris and aggregates. Post-sort analysis indicated a purity of **>**95% for GFP⁺/PI⁺ nuclei. Sorted nuclei were submitted to the University of Michigan advanced genomics core for library preparation and 10X sequencing.

### Chemogenetic neuromodulation

To investigate the role of entero-pancreatic nitrergic neurons in glucose metabolism and Ca^2+^ dynamics, a cohort of 10 *nNos*-Cre mice was used. Five mice received pancreatic intraductal infusion of AAVrg-DIO-hSyn-hM3D(Gq)-mCherry to express excitatory DREADDs in pancreas-projecting nitrergic neurons, while five received a control virus (AAVrg-DIO-hSyn-mCherry). A series of glucose tolerance tests (GTTs) were conducted before and after viral delivery (See experimental timeline in Fig. 6a).

Following completion of metabolic testing, all mice received intraperitoneal injection of two custom AAVs: AAV8-Ins-GCaMP6 (VectorBuilder) to target Ca^2+^ indicator expression to pancreatic beta cells, and AAV-PHP.S-DIO-CAG-GCaMP6 (VectorBuilder) to label peripheral nitridergic neurons. Two weeks post-injection, animals were subjected to Ca^2+^ imaging experiments.

On the day of imaging, intravital Ca^2+^ imaging was performed on the exteriorized gut and pancreas under anesthesia. Mice were then euthanized via cervical dislocation, and the pancreas was infused with agarose, sectioned into live slices, and imaged for Ca^2+^ dynamics. Slices were subsequently processed for histology.

### Intravital Ca^2+^ imaging of the pancreas and duodenum

Isoflurane (1.5-2%) anesthetized mice were fixed in the supine position on a heated imaging platform. Animal temperature was maintained at 36C. The pancreatic head and proximal duodenum were partially exteriorized through the midline abdominal incision and stabilized in a custom 3D-printed vacuum window, which eliminates movement artifacts.^38,39^ Imaging window was positioned to capture both duodenum and the pancreas for simultaneous imaging. Exteriorized tissue was imaged on a Nikon upright confocal microscope with resonant scanner and Piezo-motorized 25×/1.1NA immersion objective. Two regions of interest (ROIs) were selected: one focused on the pancreatic islet and the second one on the duodenal myenteric plexus. The automated translational stage allowed simultaneous imaging of two ROIs in in sequential XYZT mode with the temporal resolution of 2-5 seconds (depending on the distance between ROIs), and spatial resolution of 512 × 512 pixels. GCaMP fluorescence was recorded at 488Ex/510-550Em, DREADD-mCherry fluorescence was recorder at 568Ex/600-640Em. Isoflurane anesthesia induced hyperglycemia ranging 250-400 mg/dl. In all animals, pancreatic islets were active at baseline with synchronous Ca^2+^ pulses observed every 2-5 minutes. We recorded changes in GCaMP fluorescence in pancreatic beta cells and myenteric nitridergic neurons at baseline and upon IP injection of CNO (5mg/kg).

### Preparation and imaging of living pancreatic slices

Acute pancreatic slices were prepared, as previously described.^31,39^ After euthanasia, the abdomen was exposed and the pancreas infused through the common bile duct with 1.2% low gelling temperature agarose (39346-81-1, Sigma) dissolved in physiological buffer without BSA. After injection, the pancreas was extracted, cut into pieces, further embedded in agarose, and allowed to solidify at 4°C for 10 min. Pancreatic slices were cut on a vibratome (VT1000S, Leica) and incubated in physiological buffer (125 mM NaCl, 5.9 mM KCl, 2.56 mM CaCl_2_, 1 mM MgCl_2_, 25 mM HEPES, 0.1% BSA, pH 7.4) containing 7 mM glucose. Living pancreatic slices were placed in a perfusion imaging chamber (Warner Instruments) and imaged on a Leica TCS SP8 upright confocal microscope under continuous perfusion. GCaMP6 fluorescence was recorded at 488Ex/510-550Em. To identify pancreatic islets, we used the backscatter of 647 nm laser light. We recorded changes in GCaMP6 fluorescence induced by CNO (20 uM), carbachol (30 uM), and KCl (50 mM).

### Glucose tolerance test

Intraperitoneal (IPGTT) and oral glucose tolerance tests (OGTT) were conducted following a 6-hour fast. Mice received an intraperitoneal injection of the test compound—CNO (5 mg/kg), or CNO combined with nNos antagonist (ARL 17477, Tocris, 3319), or saline. Thirty minutes later, a 20% glucose solution (2 g/kg) was administered either intraperitoneally or via oral gavage. Blood glucose levels were measured at predetermined time points before and after glucose administration. Tail vein blood was collected for insulin measurement. Plasma insulin levels were measured using an ultrasensitive ELISA (Mercodia, 10-1132-01).

### Tissue collection and processing for histology

Mice were anesthetized with ketamine/xylazine (100/10 mg/kg, IP) and transcardially perfused with 4% paraformaldehyde (PFA). The stomach, intestines, pancreas, nodose ganglia (NG), dorsal root ganglia (DRG), and coeliac ganglia (CG) were harvested. Pancreata were post-fixed in 4% PFA overnight at 4 °C. NG, DRG, and CG were post-fixed for 1 h. Pancreata and ganglia were cryoprotected in 30% sucrose overnight at 4 °C, embedded in OCT, and sectioned on a Leica CM-3050S cryostat (15–30 μm for ganglia; 40–50 μm for pancreas).

### Preparation of intestinal flat mounts

After intracardial perfusion, dissected intestines and stomach were flushed with saline to remove luminal contents. Intestines were cut into 2–4 cm segments measured from the pylorus, opened longitudinally along the mesenteric border. Stomach was cut into 3 segments: fundus, corpus, and antrum. Gut segments were placed in 12 or 24 well plates where they were post-fixed in 4% PFA overnight at 4 °C. Stomach and intestinal segments were rinsed in PBS. For histological analysis we used 0.5 – 1.0 cm^2^ segments from each region. The rest of the segments were preserved in antifreeze solution at −80C. Analyzed segments were pinned mucosal side down on a Sylgard plate. The longitudinal muscle layer with attached myenteric plexus was dissected from the serosal side. The remainder of the tissue was then flipped, and the mucosa scraped to expose the submucosal plexus. These preparations (“intestinal flat mounts”) were mounted plexus-side up on microscopy slides for histological staining. Due to weak adhesion to the slide, solution exchanges during staining required extra care.

Note that successful exposure of the myenteric plexus requires careful dissection. Endogenous fluorescent neuronal markers can be identified without antibody amplification and help navigate the dissection. Frequently, if the muscle dissection is incomplete, the myenteric plexus ends up hidden in between muscle layers. In this case, it is recommended to permeabilize the tissue with 3% Triton X-100 – PBS for 1-2 h prior to blocking and extend antibody incubation times.

### Immunohistochemistry and confocal imaging

Slides with cryosections and flat mounts were blocked with Universal Blocking Reagent (Biogenex) in 0.3% Triton X-100 – PBS and incubated with primary antibodies for 48 h at room temperature or 72 hrs in 4C. After washing, Alexa Fluor-conjugated secondary antibodies (1:500 in PBS) were applied for 12–24 h. Slides were mounted with VectaShield (Vector Laboratories, H-1000). Confocal imaging was performed using a Dragonfly Spinning Disk Confocal microscope with 20× and 40× objectives (pinhole = airy 1). Images were processed in ImageJ and Imaris (Oxford Instruments). Figures show maximum projections of confocal stacks (5–30 planes, z-step = 1 μm). All primary antibodies used in this study were validated for species and application by the manufacturer and/or previously published studies.

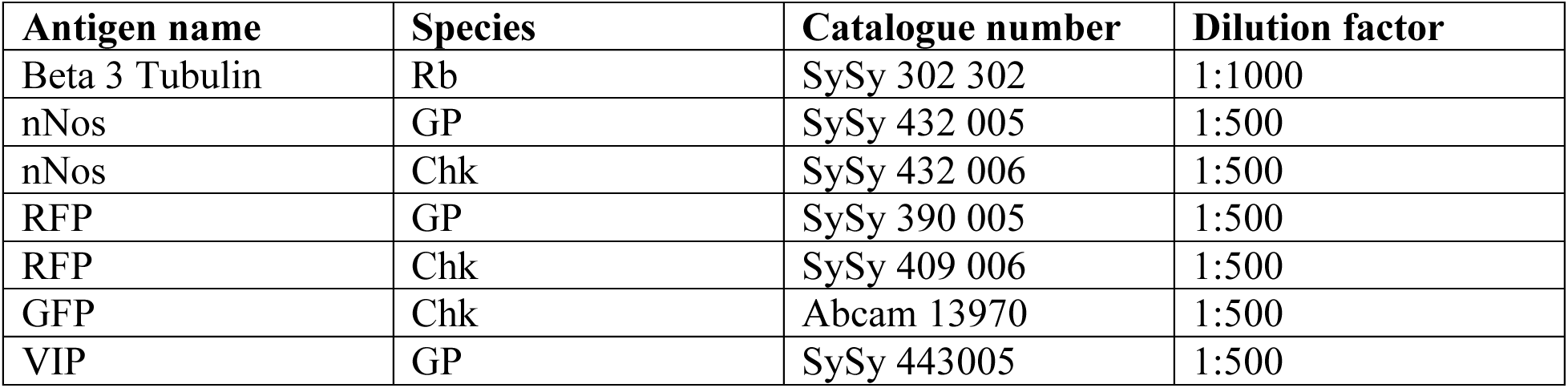

| Antigen name | Species | Catalogue number | Dilution factor |
| --- | --- | --- | --- |
| Beta 3 Tubulin | Rb | SySy 302 302 | 1:1000 |
| nNos | GP | SySy 432 005 | 1:500 |
| nNos | Chk | SySy 432 006 | 1:500 |
| RFP | GP | SySy 390 005 | 1:500 |
| RFP | Chk | SySy 409 006 | 1:500 |
| GFP | Chk | Abcam 13970 | 1:500 |
| VIP | GP | SySy 443005 | 1:500 |

### Quantification of Traced Neurons in the Myenteric Plexus and Pancreas

Retrogradely labeled neurons were manually counted in confocal image planes using ImageJ. Neuronal immunoreactivity was quantified from 3 to 15 randomly selected areas per anatomical region (each containing 100–1000 neurons), across the entire gut in 3 mice. The number of traced neurons was normalized to the total number of βIII-tubulin–positive neurons within the same field of view, and data are expressed as percentages. To estimate the absolute number of traced nitrergic neurons in the pancreas and proximal duodenum, these percentages were multiplied by published estimates of total neuronal counts in each tissue, approximately 61,000 myenteric neurons in the proximal duodenum and 2,000 neurons in the pancreas.^58–60^

### Quantification of cytosolic Ca^2+^ levels

To quantify changes in intracellular Ca^2+^ levels, we manually selected regions of interest (ROIs) around individual beta cells or myenteric neurons. We only included neurons that expressed either stimulus-evoked or baseline Ca^2+^ responses. Thus, ROI selection was biased towards bright and responding cells. In each ROI, we measured changes in mean GCaMP fluorescence intensity using ImageJ. For subsequent data analysis we used custom MatLab script. Changes in fluorescence intensity were expressed as percentage changes over baseline (ι1F/F_0_). The baseline was defined as the mean of the intensity values during the non-stimulatory control period of each recording. Most recorded cells exhibited Ca^2+^ signals during the non-stimulatory control period (baseline activity) as well as stimulus evoked responses. The strong baseline activity made it difficult to distinguish responders from non-responders based on amplitude and area under the curve, giving many false negatives and not adequately reflecting the data. For this reason, we performed peak analysis using MATLAB’s find peaks function. Peaks were identified based on a minimum prominence threshold of 10% ΔF/F. Maximum peak amplitude and maximum peak width were extracted for each cell during baseline and each stimulus. Summary statistics (mean, standard deviation, median, and mode) were computed across all cells for each parameter. Violin plots, heatmaps, and trace overlays were generated for visualization.

### Sequencing analysis

Raw sequencing reads were processed by the University of Michigan Advanced Genomics Core using the 10x Genomics Cell Ranger pipeline (version 7.0.0) with default parameters. Reads were aligned to the mm10-2020-A mouse reference transcriptome, which includes both exonic and intronic regions, using the STAR aligner embedded within Cell Ranger. The pipeline performed demultiplexing, barcode filtering, UMI counting, and cell calling based on modeling the UMI count distribution to distinguish true nuclei from background noise. Intronic reads were included in the count matrix to increase sensitivity. The resulting filtered dataset contained an estimated 247 nuclei.

Downstream processing was performed using Trailmaker visualization software (Parse Biosciences, version 1.4.1) applying standard quality control filters to exclude low-quality nuclei and potential doublets. Specifically, nuclei were filtered based on minimum and maximum thresholds of UMI counts, gene counts, and mitochondrial gene percentage, removing nuclei with aberrant values indicative of poor quality or multiplets. Following these quality control steps, the GFP-positive cluster consisted of 111 nuclei, which was homogeneous but too small for differential expression analysis. Using Trailmaker, gene expression dot plots were generated to profile gene expression of neuronal secretory markers, neuropeptides, receptors, and neuromodulatory channels within this population.

### Statistical analysis

Animals were randomly assigned to experimental and control groups. Data acquisition and analysis for Ca^2+^ imaging and metabolic assays were performed blinded when possible. In some surgical and imaging procedures, blinding was not possible due to the visibility of reporter fluorescence. For all experiments, *n* denotes biological replicates (independent animals), with numbers specified in figure legends. Sample sizes were based on prior comparable studies and deemed sufficient; no formal power calculations were performed. For Ca^2+^ imaging, peak amplitude and width during each stimulation were plotted as violin plots with median values per mouse overlaid; baseline vs. CNO intervals were compared using the Wilcoxon signed-rank test (Fig. 5g,h). For metabolic tests, AUCs were compared using unpaired Student’s t-tests (Fig. 6d,g,j), and insulin secretion data by two-way ANOVA with multiple comparisons (Fig. 6e,h,k). Two-sided tests were applied, with *P* < 0.05 considered significant. Parametric test assumptions were not formally tested; non-parametric alternatives were used when appropriate.

### Data and code availability

The RNA-seq dataset generated in this study has been deposited in NIH SRA repository under accession number SUB15538407. All other data supporting the findings of this study are available within the article and its Supplementary Information files or from the corresponding author upon reasonable request.

Custom MATLAB script used for Ca^2+^ imaging analysis includes functions for ΔF/F₀ calculation and peak analysis and is available from the corresponding author upon request. Sequencing data were processed with Cell Ranger (v7.0.0) using the embedded STAR aligner, and downstream analysis was performed in Trailmaker (Parse Biosciences v1.4.1). Imaging data were analyzed with ImageJ/Fiji (version 1.54p), Imaris (Oxford Instruments, version 10), and statistical analyses were performed in MATLAB (version R2024b) and GraphPad Prism (version 10).

### Hazards

This study did not involve agents, pathogens, or technologies subject to dual-use research of concern regulations. No unusual hazards were encountered during the course of this research.

## Acknowledgements

The authors thank members of the Randy Seeley and Martin Myers laboratories at the University of Michigan for their scientific input, technical support, and the stimulating research environment. We are especially grateful to Dr. Alfor Lewis for performing surgeries and providing surgical training, Dr. Chelsea Hutch for conducting glucose tolerance tests and technical assistance, and Abigail Tomlinson for her help and training in nuclei isolation for RNA sequencing. This work was funded by NIH grants R56DK084321 (A.C.), R01DK084321 (A.C.), R01DK111538 (A.C.), R01DK113093 (A.C.), U01DK120456 (A.C.), U01DK135017 (A.C), R33ES025673 (A.C.), R21ES025673 (A.C.), R01DK130328 (A.C.), and R01DK138471 (A.C); P30DK089503 (R.J.S.), R01DK133140 (R.J.S.); M.S. received a postdoctoral fellowship F32DK132795 (M.S.);

## Contributions

Conceptualization: M.S, R.J.S., and A.C. Data curation: M.S. Formal analysis: M.S., R.J.S., and A.C. Funding acquisition: M.S, R.J.S., and A.C. Investigation: M.S., A.T., N.L. Methodology: M.S., A.T. Project administration: M.S, R.J.S., and A.C. Resources: M.S, R.J.S., and A.C. Software: M.S. Supervision: M.S, R.J.S., and A.C. Validation: M.S. Visualization: M.S. and A.C. Writing, original draft: M.S., R.J.S., and A.C. Writing, reviewing and editing: all authors.

## Ethics declarations

RJS has received research support from Fractyl, AstraZeneca, Congruence Therapeutics, Eli Lilly, Diasome and Amgen. RJS has been a paid consultant for Eli Lilly, CinRx, Crinetics, Amgen, Gallant and Nuanced Health. RJS has equity in Nuanced Health, Coro Bio, Eccogene, Fractyl and Rewind.

