## Extended data for "A neural entero-pancreatic pathway that regulates insulin secretion and glucose tolerance"

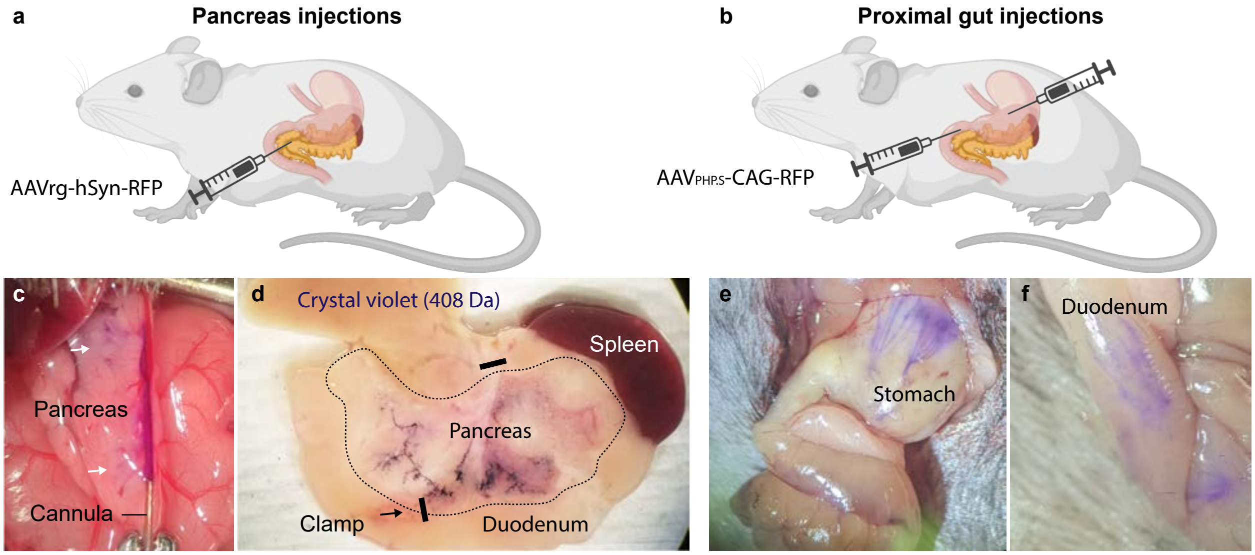


**Extended Data Figure 1. Viral injection approaches illustrated with Crystal Violet dye.**

**a, c, d**: Intraductal pancreatic infusion for viral delivery to the pancreas. Figure adapted from Makhmutova *et al*., 2021[31] (technique described in Xiao *et al*., 2014[56]).

**b, e, f**: Submuscular injections into the stomach and proximal duodenum, as described in Han *et al*., 2018 [57].

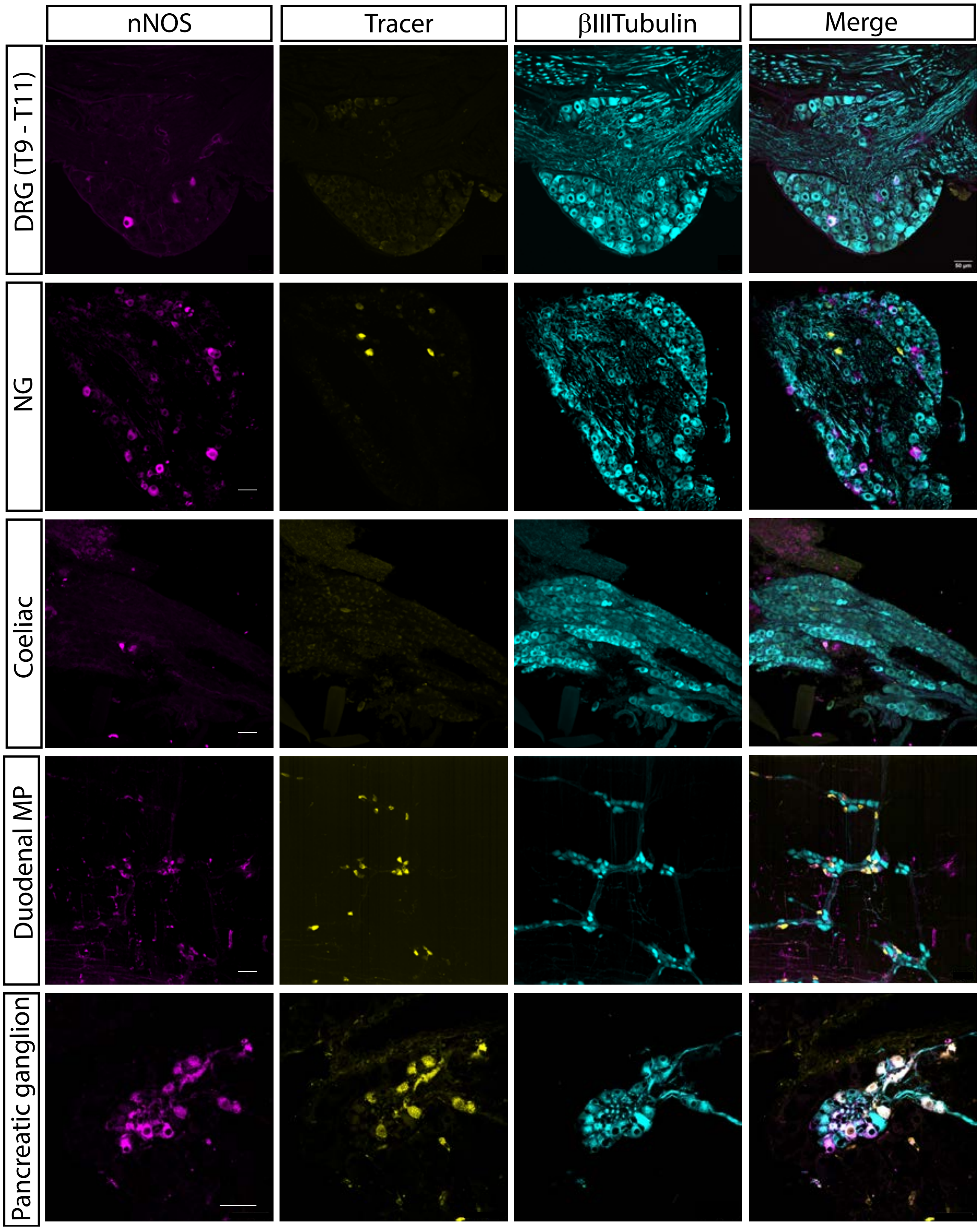


**Extended Data Figure 2. Representative confocal images showing immunostaining for neuronal nitric oxide synthase.**

(nNOS, magenta), retrograde tracer AAVrg-hSyn-mCherry (yellow), and βIII-tubulin (cyan) in dorsal root ganglia (DRG, T9–T11), nodose ganglia (NG), coeliac ganglia, duodenal myenteric plexus (MP), and pancreatic ganglia. Merged images reveal colocalization of tracer and nNOS in myenteric and pancreatic ganglia, absence of colocalization in the nodose ganglion, and no tracer labeling in coeliac ganglion or DRG. Scale bars, 50 μm.

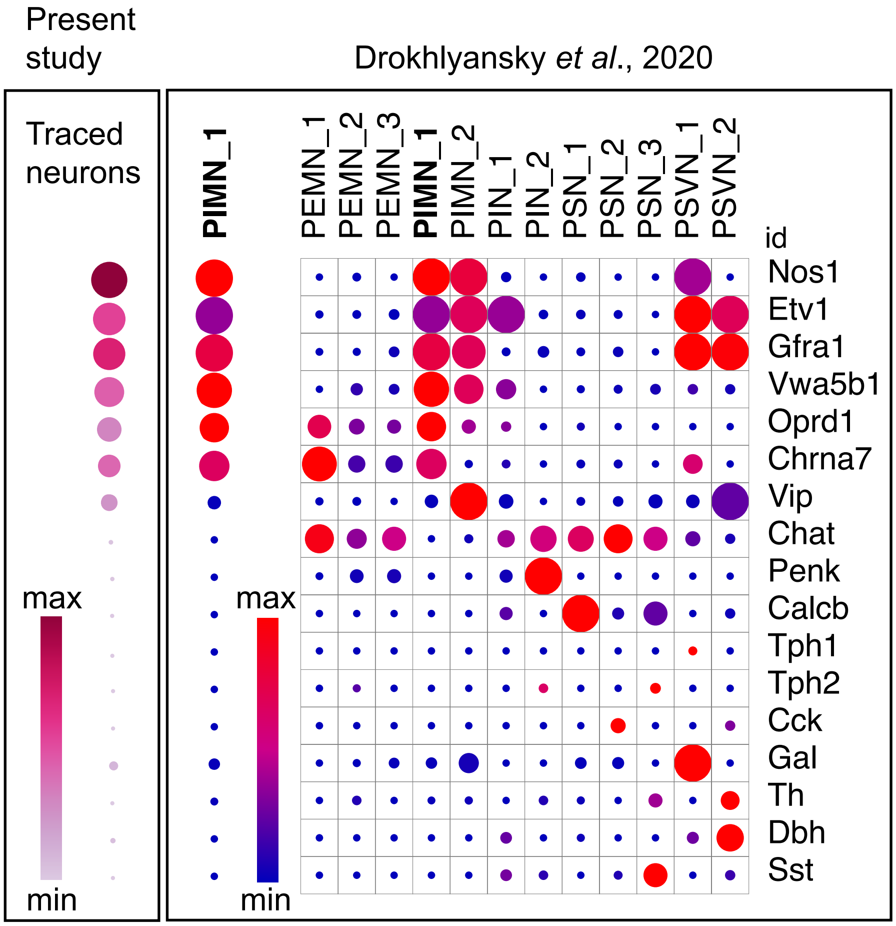


**Extended Figure 3. Comparison of gene expression in traced entero-pancreatic myenteric neurons with small intestinal neuronal clusters from Drokhlyanski *et al.*, 2020.**

Dot plots showing gene expression profiles, where dot size represents the fraction of nuclei expressing the gene and dot color indicates mean expression level in expressing nuclei. The left column depicts the signature gene expression profile of entero-pancreatic neurons traced in the present study. The right panel shows gene expression across myenteric neuronal clusters (PIMN, PEMN, PIN, PSN) assigned to specific enteric nervous system subtypes in Drokhlyanski *et al.* The traced entero-pancreatic neurons exhibit a gene expression signature closely matching the PIMN1 cluster of primary inhibitory motor neurons cluster 1.

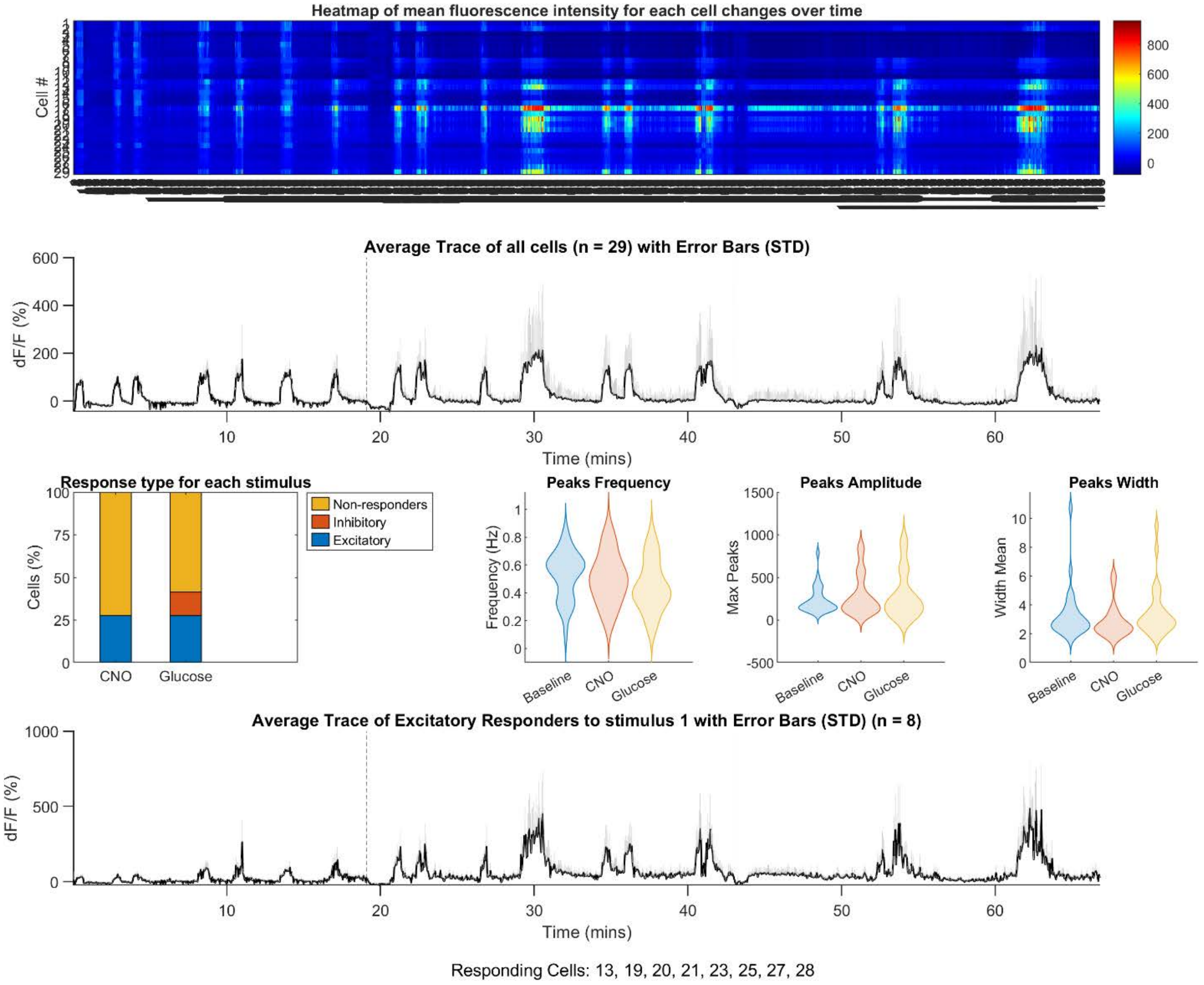


**Extended Figure 4. Intravital $Ca^{2+}$ imaging of pancreatic islet activity in vivo – Mouse 1**

nNos-Cre mice received intraductal pancreatic injections of AAVrg-DIO-hM3D(Gq)-mCherry to express the excitatory DREADD receptor in entero-pancreatic neurons and intraperitoneal injections of AAV8-Ins-GCaMP6 to express the $Ca^{2+}$ indicator GCaMP6 in pancreatic beta cells. $Ca^{2+}$ imaging was performed in an anesthetized mouse. The upper panel shows a heat map of *in vivo* $Ca^{2+}$ responses, with each horizontal row representing an individual beta cell and color intensity corresponding to changes in mean fluorescence ($\Delta F/F_0$). Traces below show normalized $\Delta F/F$ from individual beta cells over time. CNO application, indicated by a dashed line, elicited synchronized $Ca^{2+}$ transients across beta cells compared to baseline. Violin plots display maximum peak frequency, amplitude, and width for each cell. The bottom panel shows the average trace of responding beta cells.

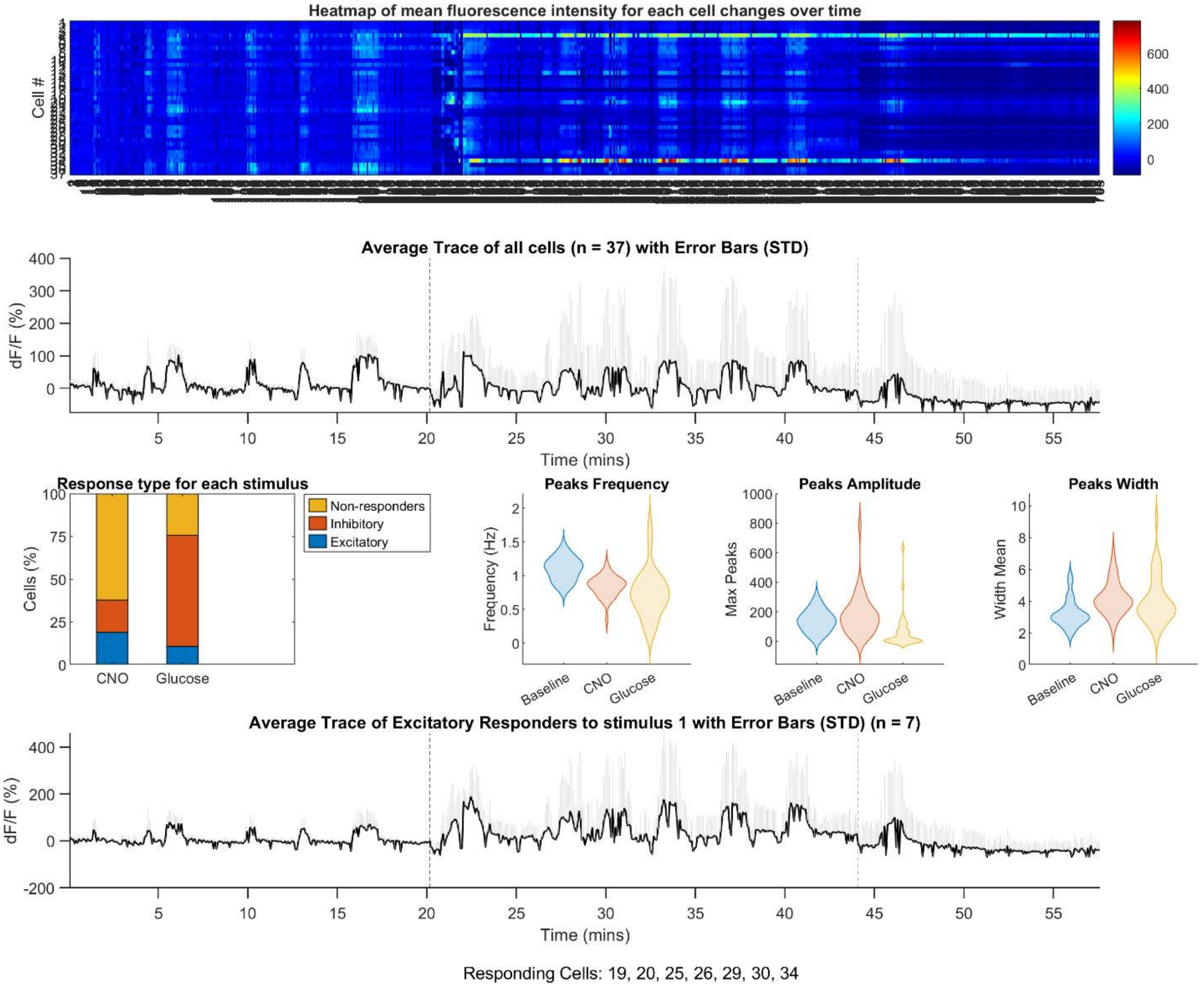


**Extended Figure 5. Intravital $Ca^{2+}$ imaging of pancreatic islet activity in vivo – Mouse 2**

nNos-Cre mice received intraductal pancreatic injections of AAVrg-DIO-hM3D(Gq)-mCherry to express the excitatory DREADD receptor in entero-pancreatic neurons and intraperitoneal injections of AAV8-Ins-GCaMP6 to express the $Ca^{2+}$ indicator GCaMP6 in pancreatic beta cells. $Ca^{2+}$ imaging was performed in an anesthetized mouse. The upper panel shows a heat map of *in vivo* $Ca^{2+}$ responses, with each horizontal row representing an individual beta cell and color intensity corresponding to changes in mean fluorescence ($\Delta F/F_0$). Traces below show normalized $\Delta F/F$ from individual beta cells over time. CNO application, indicated by a dashed line, elicited synchronized $Ca^{2+}$ transients across beta cells compared to baseline. Violin plots display maximum peak frequency, amplitude, and width for each cell. The bottom panel shows the average trace of responding beta cells.

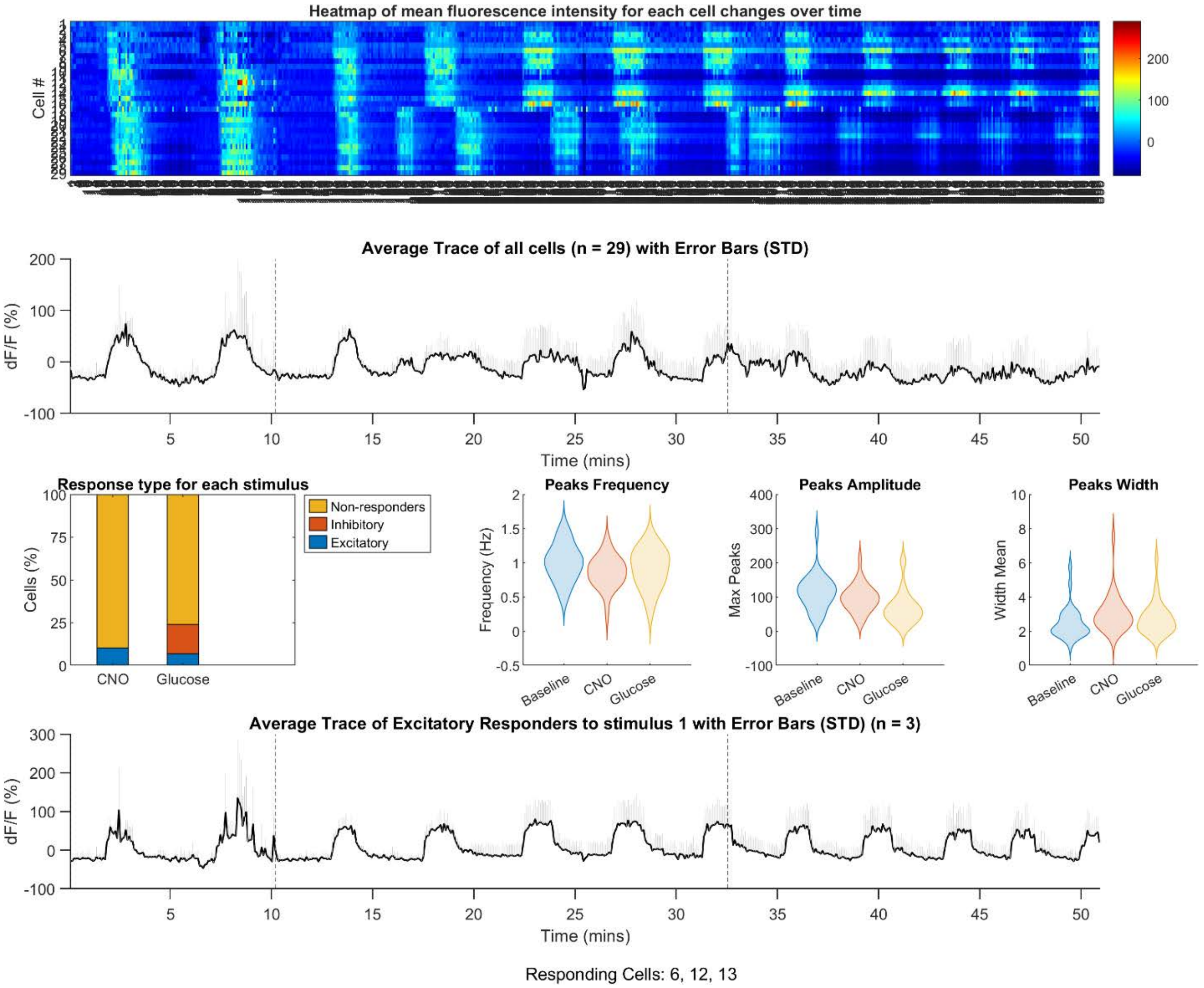


**Extended Figure 6. Intravital $Ca^{2+}$ imaging of pancreatic islet activity in vivo – Mouse 3**

nNos-Cre mice received intraductal pancreatic injections of AAVrg-DIO-hM3D(Gq)-mCherry to express the excitatory DREADD receptor in entero-pancreatic neurons and intraperitoneal injections of AAV8-Ins-GCaMP6 to express the $Ca^{2+}$ indicator GCaMP6 in pancreatic beta cells. $Ca^{2+}$ imaging was performed in an anesthetized mouse. The upper panel shows a heat map of *in vivo* $Ca^{2+}$ responses, with each horizontal row representing an individual beta cell and color intensity corresponding to changes in mean fluorescence ($\Delta F/F_0$). Traces below show normalized $\Delta F/F$ from individual beta cells over time. CNO application, indicated by a dashed line, elicited synchronized $Ca^{2+}$ transients across beta cells compared to baseline. Violin plots display maximum peak frequency, amplitude, and width for each cell. The bottom panel shows the average trace of responding beta cells.

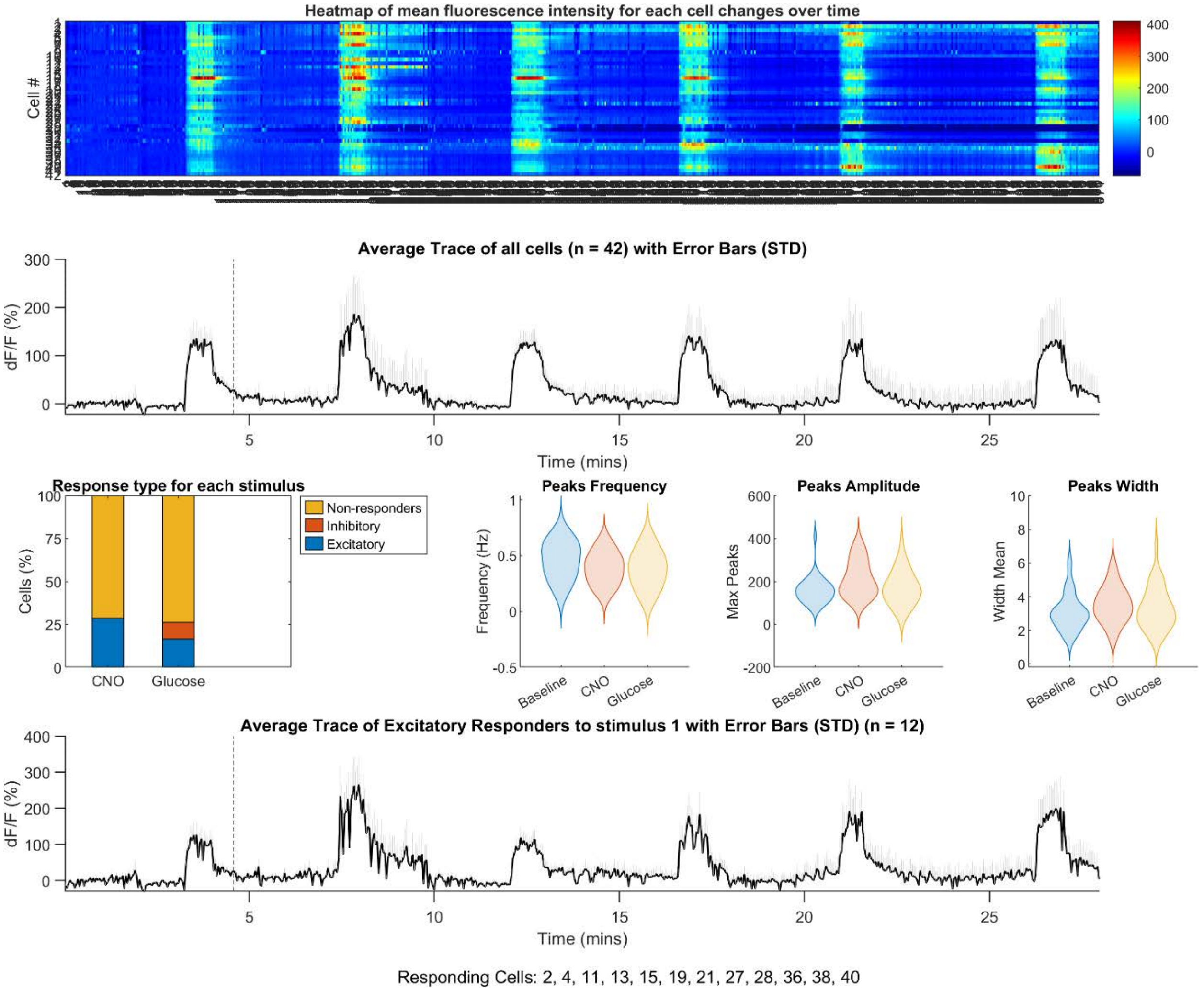


**Extended Figure 7. Intravital $Ca^{2+}$ imaging of pancreatic islet activity in vivo – Mouse 1**

nNos-Cre mice received intraductal pancreatic injections of AAVrg-DIO-hM3D(Gq)-mCherry to express the excitatory DREADD receptor in entero-pancreatic neurons and intraperitoneal injections of AAV8-Ins-GCaMP6 to express the $Ca^{2+}$ indicator GCaMP6 in pancreatic beta cells. $Ca^{2+}$ imaging was performed in an anesthetized mouse. The upper panel shows a heat map of *in vivo* $Ca^{2+}$ responses, with each horizontal row representing an individual beta cell and color intensity corresponding to changes in mean fluorescence ($\Delta F/F_0$). Traces below show normalized $\Delta F/F$ from individual beta cells over time. CNO application, indicated by a dashed line, elicited synchronized $Ca^{2+}$ transients across beta cells compared to baseline. Violin plots display maximum peak frequency, amplitude, and width for each cell. The bottom panel shows the average trace of responding beta cells.

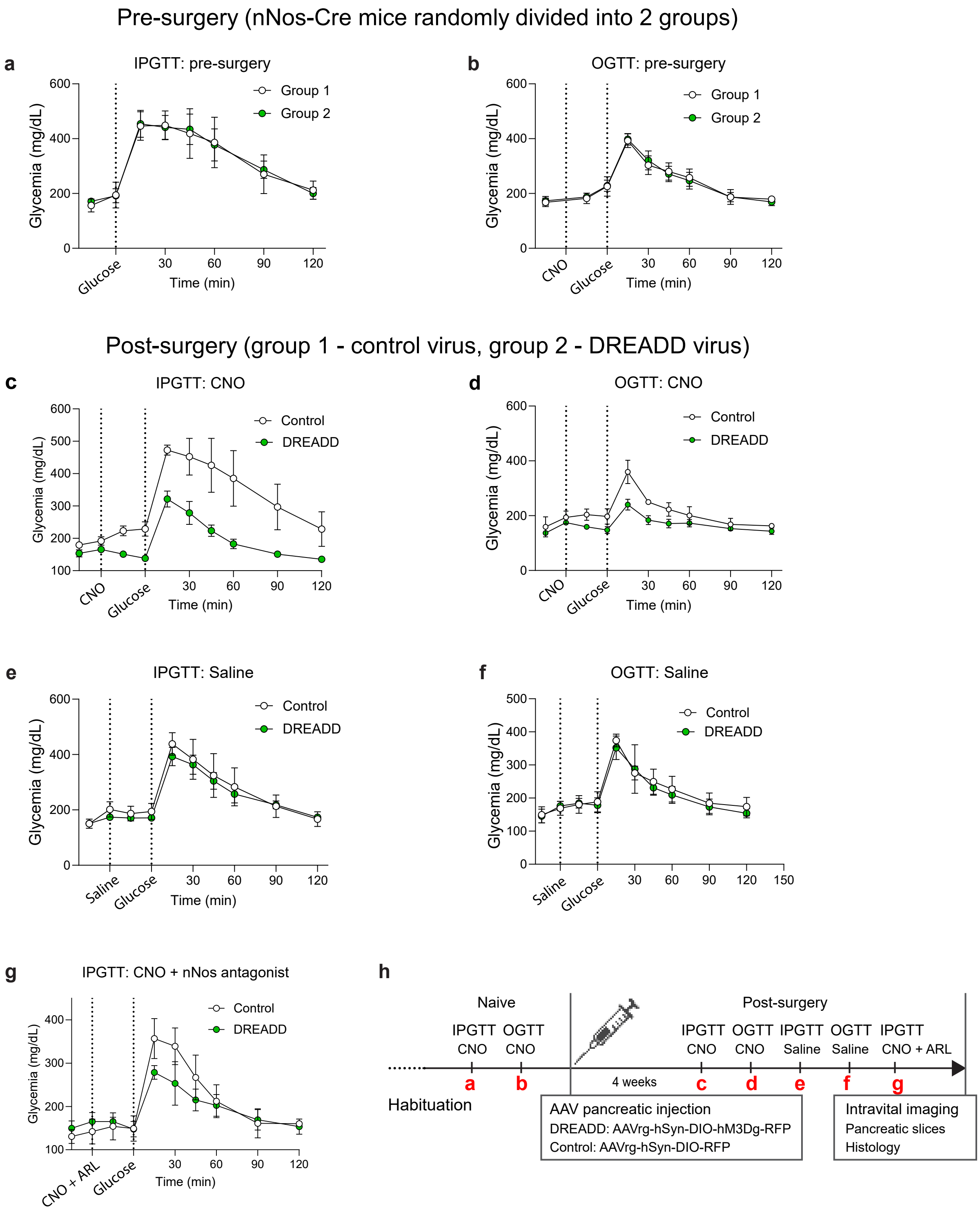


**Extended Figure 8. Glucose tolerance in *nNos*-Cre mice before and after DREADD activation of entero-pancreatic neurons.**

**a-b**, Pre-surgery intraperitoneal (IPGTT; a) and oral (OGTT; b) glucose tolerance tests in mice assigned to control or DREADD groups.

**c-d**, Post-surgery IPGTT (c) and OGTT (d) following CNO administration (5 mg/kg).
**e-f**, Post-surgery IPGTT (e) and OGTT (f) following saline administration.
**g**, Post-surgery IPGTT following CNO with nNOS antagonist (ARL17477, 10 mg/kg).
**h**, Experimental timeline. Data are mean ± s.e.m, n = 5 mice per group

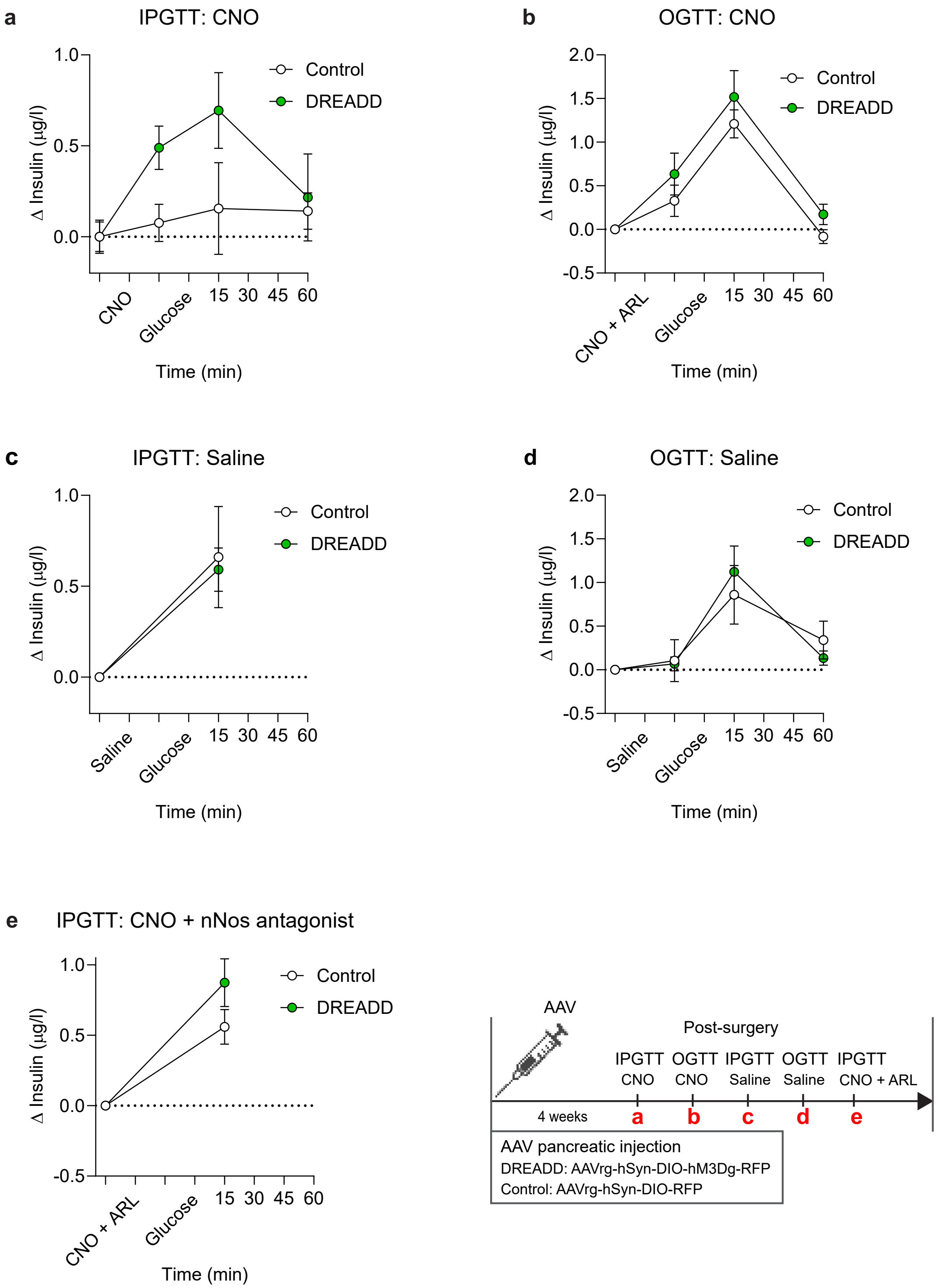


**Extended Figure 9. Insulin secretion during glucose tolerance tests in *nNos*-Cre mice after DREADD activation of entero-pancreatic neurons.**

**a-b**, Insulin secretion during post-surgery intraperitoneal (IPGTT; a) and oral (OGTT; b) glucose tolerance tests following CNO administration (5 mg/kg).

**c-d**, Insulin secretion during Post-surgery IPGTT (c) and OGTT (d) following saline administration.
**e**, Insulin secretion during post-surgery IPGTT following CNO (5 mg/kg) with nNOS antagonist (ARL17477, 10 mg/kg).
Inset: experimental timeline. Data are mean ± s.e.m, n = 5 mice per group
