## Supplementary data for "A neural entero-pancreatic pathway that regulates insulin secretion and glucose tolerance"

**Video 1. Terminals of enteric neurons in the pancreas.**

3D reconstruction of a pancreatic islet from C57BL/6 mice injected with AAVrg-hSyn-mCherry into the gut wall to trace enteric neuron terminals projecting to the pancreas. Pancreatic sections were immunostained for βIII-tubulin (cyan), vasoactive intestinal peptide (VIP, magenta), mCherry (yellow), and DAPI (blue). Surfaces were rendered in Imaris, revealing mCherry$^+$ varicosities co-stained with VIP within the islet. Some varicosities penetrate the islet parenchyma and contact βIII-tubulin$^+$ pancreatic neurons.

**Video 2.** In vivo recording of $Ca^{2+}$ activity in pancreatic beta cells at baseline hyperglycemic conditions (> 250 mg/dl) and in response to intraperitoneal CNO injection (5 mg/kg).

**Video 3.** In vitro recording of $Ca^{2+}$ activity in pancreatic beta cells at baseline 7mM glucose and in response to perfusion with CNO (20 mM), Carbachol (CCh, 30 mM), and KCl (50 mM).
